# A triple serine motif in the intracellular domains of sortilin-related receptors SorCS1-3 regulates neurotrophic activity

**DOI:** 10.1101/2024.10.03.616089

**Authors:** Anders Dalby, Mathias Kaas, Lars Meinertz-Byg, Signe Bundgaard Christiansen, Simon Bøggild, Per Qvist, Jens R. Nyengaard, Peder Madsen, Simon Mølgaard, Simon Glerup

## Abstract

The Vps10p-domain receptors SorCS1-3 have been repeatedly associated with the development of neurological and psychiatric disorders. They have emerged as key regulators of synaptic activity and neurotrophic signaling, but the underlying molecular mechanism remains poorly understood. Here we report that the SorCS1-3 intracellular domains (ICDs) contain a conserved triple serine motif that potentially functions as a signaling switch to induce neurotrophic signaling in hippocampal neurons. We demonstrate that phosphorylation mimicking mutations of the SorCS1-3 triple serine motifs display neurotrophic activity independently of both their extracellular domains (ECDs) and BDNF, and that the substitution of serines to alanines renders neurons less responsive to BDNF. Hence, we develop triple serine motif-based cell-penetrating peptides that modulate downstream signaling kinases of the BDNF pathway, ultimately activating the transcription factor CREB. Taken together, we provide the first mechanistic insights into SorCS1-3 mediated neurotrophic signaling and use this knowledge to develop pharmacologically active modulators.

## Introduction

SorCS1, SorCS2 and SorCS3 belong to the Vps10p-domain receptor family also encompassing sortilin and SorLA. GWAS, exome sequencing, linkage analysis and copy number variation studies have repeatedly implicated *SORCS1-3* in several neurological and psychiatric disorders, but the molecular mechanisms underlying these associations remain unknown [1–11]. The extracellular domains (ECDs) of SorCS1-3 share the N-terminal Vps10p domain, followed by a leucine-rich domain. The ECDs of SorCS1-3 bind to a variety of ligands, including pro and mature neurotrophins of nerve growth factor (NGF), brain-derived neurotrophic factor (BDNF) and progranulin as well as TrkB, p75NTR and NMDAR subunits [12–15]. SorCS1-3 are central for the proper function of these ligands as they influence their subcellular processing or act as co-receptors for signal transduction.

Among SorCS1-3, the biological role of SorCS2 is the best characterized. SorCS2 regulates signals involved in axonal retraction and growth cone collapse through proBDNF and proNGF [16, 17], while also regulating trafficking of the BDNF receptor, TrkB, to the postsynaptic membrane to induce synaptic plasticity in the hippocampus [14]. Additionally, SorCS2 has been shown to direct the surface expression of both NMDA receptor subunits and neuronal amino acid transporter EAAT3, illustrating the multifaceted roles of SorCS2 as a regulator of pro-neurotrophin and neurotrophin responses in neurons as well as postsynaptic receptor composition [15, 18, 19]. In line with this, *Sorcs2* KO mice display impaired spatial, contextual and social memory, enhanced neuronal cell death, increased mortality during epileptic seizures and several neuropsychiatric phenotypes, including hyperactivity, altered response to stimulants and impaired prepulse inhibition [14–16, 18–20].

All Vps10p-domain receptors have short intracellular domains (ICDs) containing putative binding motifs for several adaptor proteins, which for SorCS1-3 include AP-2, PSD95 and PICK1 [21–23]. The unique combination of such motifs across SorLA, sortilin and the SorCS1-3 receptors enable distinct trafficking routes for each receptor [24]. SorCS1-3 are preferentially involved with Rab11 positive endosomal trafficking important for recycling of receptors including neurexins, TrkB and neurotransmitter receptor subtypes, potentially via dynein and kinesin adaptor proteins [15, 18, 19, 23, 25–27]. In contrast, sortilin and SorLA sort ligands to lysosomal compartments and early endosomes, respectively [28–30]. This demonstrates the diverse functions mediated by Vps10p-domain receptors through both ECD ligand specificity and differential subcellular sorting abilities of their ICDs. Apart from the critical role of the SorCS2-ICD in BDNF-induced Long-Term Potentiation (LTP) and neuronal branching [14], its cellular functions remain largely unknown.

Here, we studied the role of the SorCS2-ICD in BDNF/TrkB mediated signaling. We show that the effects of the SorCS2-ICD on downstream BDNF signaling are mediated by a conserved triple serine motif, and possibly regulated by phosphorylation. In addition, we demonstrate that phosphomimetics of the SorCS2-ICD can mediate neurotrophic signaling independently of both the SorCS2 ECD and BDNF itself. This activity is also conserved in the SorCS1- and SorCS3-ICDs. Utilizing this knowledge, we developed peptide phosphomimetics of the SorCS2-ICD that induce activation of distinct signaling cascades downstream of SorCS2/BDNF/TrkB, leading to increased dendritic branching and PSD-95 clustering. More specifically, using phospho-kinase arrays, we observe that the SorCS2-ICD derived phosphomimetic peptides modulate the MAPK pathway, ultimately leading to phosphorylation of the transcription factor CREB. Lastly, the ICD phosphomimetics induced transcriptomic changes consistent with activation of Jun and c-Fos further substantiating the engagement with neurotrophic signaling. Taken together, our data uncover a novel function of the SorCS1-3 ICDs in mediating neurotrophic signaling as well as the potential for using SorCS-ICD mimetics for therapeutic purposes.

## Results

### SorCS2 is important for BDNF-induced MAPK pathway activation

To understand the mechanistic role of SorCS2 in BDNF/TrkB signaling, we stimulated WT and *Sorcs2* KO hippocampal neurons at DIV5 for 10 min with BDNF (1 nM) and submitted homogenates to an antibody array encompassing 877 immobilized antibodies capturing a wide variety of signaling proteins (KINEXUS). In WT neuronal homogenates, the array detected BDNF-induced phosphorylation of multiple targets known to be involved in neuronal gene transcription and protein translation in addition to synapse formation and plasticity (Fig. 1a). These included well-known proteins within the BDNF-signaling pathway: TrkB (Y702), Akt (S473), CAMK (T177, Y240), PKC (Y195, T507), STAT3 (S727), NMDAR2B (Y1474) and Src (Y530, Y419). Notably, we also observed phosphorylation of several unexpected targets outside the classical BDNF signaling pathway (Fig. 1b), such as GSK3a (S21), PKCd (Y334), ErbB4 (Y875, Y733), PTPN21 (S637) and p53 (S315). We found strongly decreased activation of these targets in *Sorcs2* KO neurons, suggesting that SorCS2 plays a critical role in the transduction of multiple BDNF-induced signaling cascades. KEGG pathway enrichment analysis of the phospho-array data further identified the MAPK signaling pathway among differentially phosphorylated proteins in SorCS2 KO neurons versus WT neurons in response to BDNF (Fig. S1a).

**Fig. 1:**
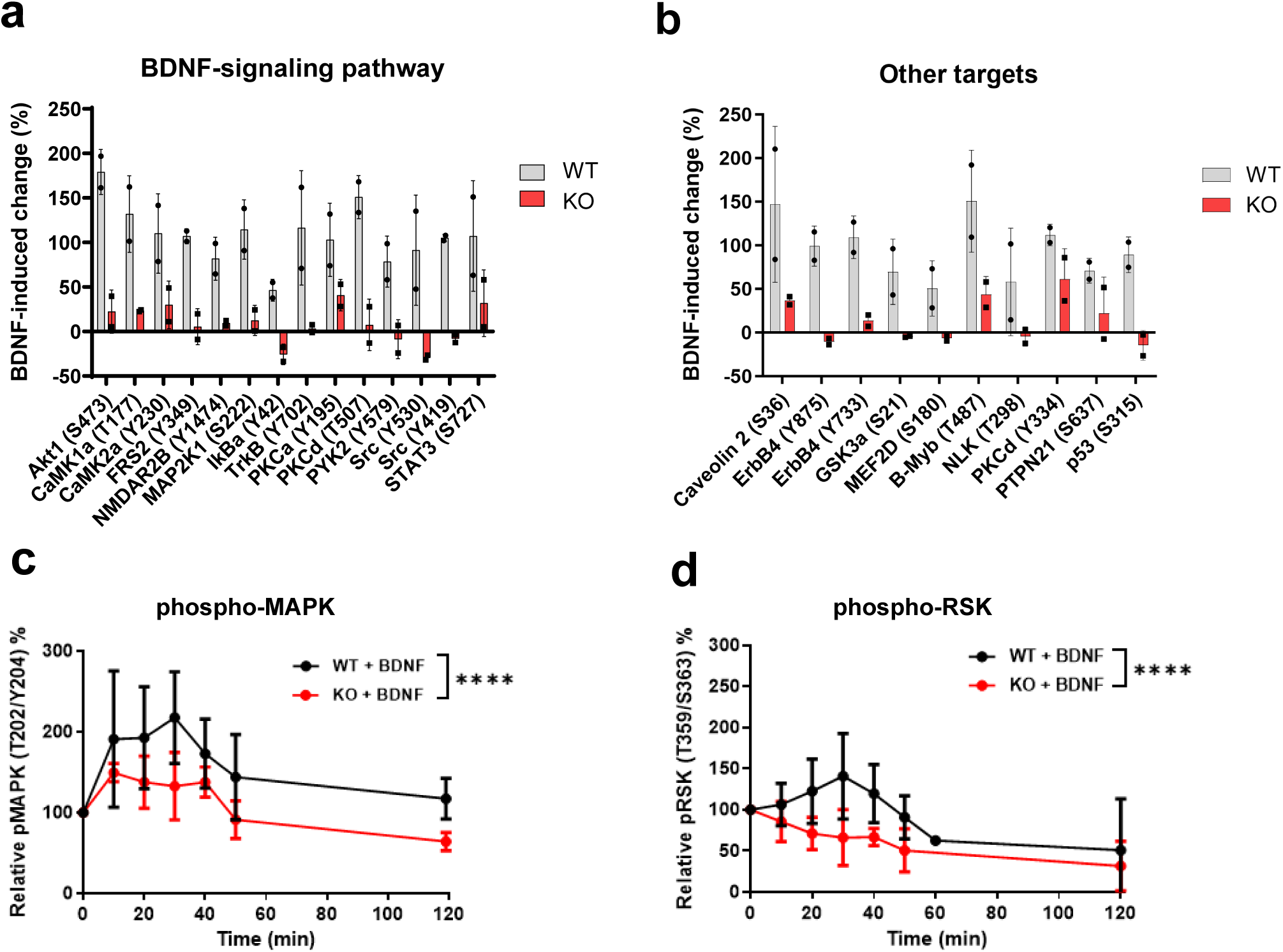
Lack of SorCS2 hampers BDNF-signaling. **(a)** Identification of downstream targets in the BDNF-signaling cascade and **(b)** targets of other pathways using a phospho-antibody array in hippocampal neurons (DIV5) from WT and KO mice stimulated with or without BDNF (1 nM, 10 min, n=2). Hits shown with p<0.05 and a percentage-change of +-45%. **(c-d)** Time course of BDNF-induced activation of MAPK and RSK in WT and KO hippocampal neurons (n=5 of each genotype). Significance was calculated using ordinary two-way ANOVA of main effects, ****p < 0.0001. Data is represented as mean ± SD.

The effect of BDNF on synaptic protein expression is known to require sustained activation of MAPK. This leads to the phosphorylation of the kinase RSK and its translocation to the nucleus, which ultimately phosphorylates CREB, allowing it to induce gene transcription and neurotrophic support [31]. We therefore studied the time course of MAPK and RSK phosphorylation in hippocampal neurons from WT and KO stimulated with BDNF and found their phosphorylation to be significantly reduced in the absence of SorCS2 (Fig. 1c-d and Fig. S1b-c). Importantly, there were no differences in the baseline phosphorylation signal, in agreement with previous studies in *Sorcs2* KO mice (Fig S1d-e) [18].

### The ICD of SorCS2 contains a conserved triple serine motif required for neurotrophic signaling

Previous studies from our group have demonstrated that the ICD of SorCS2 is critical for BDNF-mediated effects on the induction of LTP and dendritic branching, and that the BDNF response in *Sorcs2* KO neurons can be rescued by transfection with plasmids encoding full length SorCS2 (FL SorCS2) but not a SorCS2 variant lacking the ICD [14] (Fig. 2a). We therefore used this assay to identify the specific region of the SorCS2-ICD, which is required for maintaining the BDNF-induced neurotrophic effects. Consequently, we co-transfected *Sorcs2* KO neurons with truncated (TC) ICD variants of SorCS2 (TC1, TC2, TC3 and tailless, Fig. 2b) with GFP, while using the FL SorCS2 receptor (WT) as a positive control and measured neuronal branching of GFP positive neurons in the absence or presence of BDNF. Interestingly, removal of the C-terminal 20 amino acids (1152-1172) in the TC1 variant had a significant effect on the ability of SorCS2 to mediate BDNF-induced neuronal branching (Fig. S2a). This region is known to be critical for linking SorCS2 to kinesin and dynein motor proteins via binding to KLC1 and DYNLT3, respectively, and thereby regulating SorCS2 anterograde and retrograde transport [23]. Despite the potential importance of this region for SorCS2 activity, TC1 was still able to provide some rescue of BDNF activity in transfected neurons (Fig. 2c-d). The removal of amino acids 1138-1151 in TC2 did not change the rescue ability compared to TC1, suggesting that this sequence stretch does not play a critical role (Fig. 2c-d, Fig. S2). However, removal of amino acids 1117-1137 resulted in a completely inactive SorCS2 variant, suggesting the importance of this region. Importantly, all SorCS2 variants expressed at similar levels compared to FL SorCS2 in transfected HEK293 cells (Fig. S2b).

**Fig. 2:**
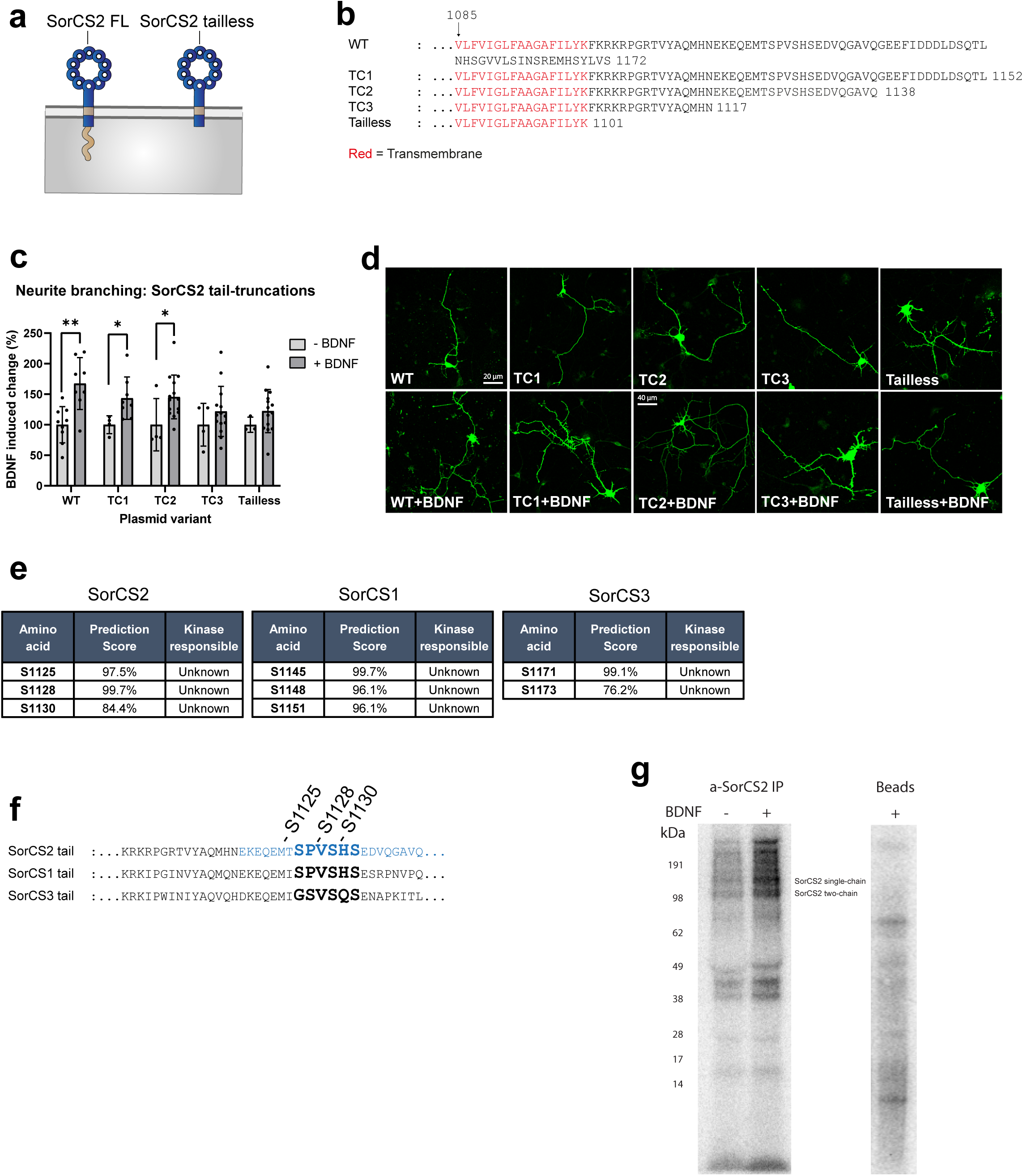
The SorCS2-tail contains a conserved serine motif phosphorylated upon BDNF-signaling. **(a)** Schematic of SorCS2 full-length (FL) receptor and tailless version. **(b)** Sequences of wildtype (WT) SorCS2-tail and truncated SorCS2-ICD variants (TC1, TC2, TC3) and tailless SorCS2. Transmembrane region is marked in red. Numbers indicate amino acid positions. **(c)** Neuronal branching in response to BDNF (1 nM) in SorCS2 KO hippocampal neurons co-transfected with GFP and truncated variants of SorCS2 receptor (n=4-14 coverslips per group with 5-25 neurons analysed per coverslip, mean branches per neuron showed) and **(d)** representative images. Significance of neurite branching experiments was calculated using pair-wise t-tests (*p<0.05, **p < 0.01). Data is represented as mean ± SD. **(e)** Prediction of chance of phosphorylation of serine motif in SorCS1, SorCS2 and SorCS3-ICD by Netphos 3.1 with indicated amino acid positions. **(f)** Sequence alignment of SorCS1, SorCS2 and SorCS3-ICDs and their serine motif. Important region in SorCS2 for BDNF-induced branching is marked in blue and position of serines in SorCS2 is highlighted. **(g)** Visualization of ^32^P-labelled proteins after pulldown with anti-SorCS2 conjugated beads (left) or unconjugated beads as negative control (right).

As intracellular signaling cascades are mainly driven by regulatory modifications involving protein phosphorylation and dephosphorylation, we assessed the SorCS2-ICD for potential phosphorylation motifs (Tyr, Ser, Thr) using the machine-learning based tool NetPhos 3.1. The algorithm identified three serines with a high prediction score for phosphorylation (>0.8), namely S1125, S1128 and S1130, which were all within the TC2 tail-fragment. This triple serine motif is conserved across species (Fig. S4a) and is also present in SorCS1 and SorCS3, albeit the SorCS3 ICD contains only two of the three serines predicted to be phosphorylated (Fig 2e-f). No other serine, tyrosine, or threonine residues, respectively, showed high confidence predictions (>0.8).

To study if SorCS2 is phosphorylated and if this could be influenced by BDNF, we labelled DIV7 hippocampal neurons with radioactive phosphate (^32^P) and treated them with BDNF (1 nM) or vehicle for 10 minutes. SorCS2 was subsequently captured from lysates by immunoprecipitation using antibodies raised against the intracellular domain. Interestingly, phosphorylated bands migrating at similar size as one-chain SorCS2, two-chain SorCS2, and the processed C-terminal fragment encompassing the juxtamembrane, transmembrane and cytoplasmic region [16], suggesting that SorCS2 is phosphorylated in both the extracellular and cytosolic parts, in accordance with phosphoproteomic databases [32, 33] (Supplementary Table 1). The intensity of bands was enhanced by the presence of BDNF, suggesting that SorCS2 phosphorylation is affected by BDNF-induced signaling (Fig. 2g). Of note, additional bands were visible in the SorCS2 precipitates, possibly indicating unknown phosphorylated proteins interacting with SorCS2.

### Phosphomimetic mutations in SorCS2 induce neurotrophic effects

Receptor Tyrosine Kinases (RTKs), including TrkB, are auto-phosphorylated upon ligand binding, which creates docking sites for signaling complexes or directly activates intracellular kinase domains to convey downstream signaling via phosphorylation of specific serine, threonine or tyrosine residues on selected targets [31]. To assess if the triple serine motif in the SorCS2-ICD is important for the transmission of BDNF/TrkB-induced downstream neurotrophic signaling, we first constructed FL SorCS2 variants by single serine-to-alanine substitutions of S1125, S1128 and S1130 as well as a complete mutated triple serine motif S1125A/S1128A/S1130A (denoted S3A) mutant (Fig. 3a) and assessed their ability to rescue BDNF-induced neuronal branching in S*orcs2* KO neurons. We observed a loss of BDNF response on dendritic branching for S3A, S1128A and S1130A variants of the SorCS2 FL receptor, while the S1125A variant maintained a partial response (Fig. 3b-c and S4b). No clear change in subcellular localization was observed for either of the variants in HEK293 cells (Fig. S5), suggesting that the loss of neurotrophic signaling is not due to aberrant trafficking of SorCS2.

**Fig. 3:**
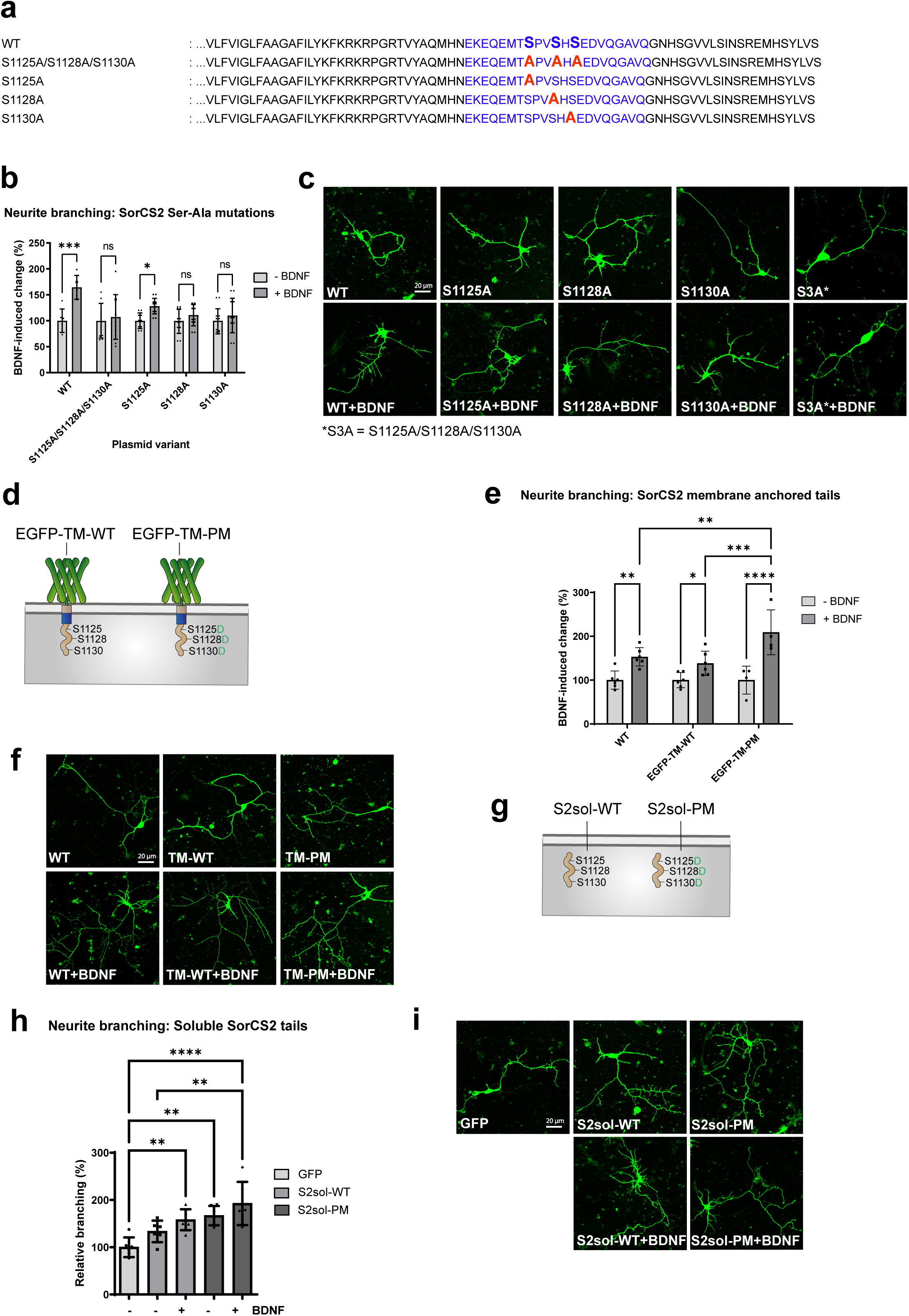
Soluble phospho-mimetic SorCS2-ICDs induces neuronal branching independent of BDNF. **(a)** Sequences of wildtype (WT) SorCS2-ICD and variants with serine-to-alanine mutations (S1125A, S1128A, S1130A, S1125A/S1128A/S1130A). Important region in SorCS2 for BDNF-induced branching is marked in blue and serine-to-alanine mutations in SorCS2 is highlighted in red. **(b)** Neurite branching in response to BDNF (1 nM) in *Sorcs2* KO hippocampal neurons transfected with GFP and SorCS2 FL (WT) or serine-to-alanine mutation variants (n=6-12 coverslips per group with 5-15 neurons analysed per coverslip) and **(c)** representative images. Significance of neurite branching experiments was calculated using two-way ANOVA (*p < 0.05, and ***p =0.0001). **(d)** Drawing of SorCS2-transmembrane (TM) and tail linked to EGFP-construct with either wild-type serines (EGFP-TM-WT) or aspartic acid substitutions (EGFP-TM-PM) as a phospho-mimetic of serine motif. **(e)** Neuronal branching in response to BDNF (1 nM) in *Sorcs2* KO hippocampal neurons transfected with GFP and SorCS2-ICD constructs linked to EGFP (n=4-6 coverslips per group with 5-15 neurons analysed per coverslip) and **(f)** representative images. Significance of neurite branching experiments was calculated using two-way ANOVA (*p < 0.05, **p<0.01, ***p =0.0006 and ****p<0.0001). **(g)** Drawing of soluble SorCS2 tails with aspartic acid substitutions indicated. **(h)** Neurite branching of *Sorcs2* KO hippocampal neurons transfected with GFP and soluble SorCS2-ICD constructs (WT or phosphomimetic aspartic acids) in combination with 1 nM BDNF (n=6 coverslips per group with 5-15 neurons analysed per coverslip) and **(i)** representative images. Data is represented as mean ± SD.

We subsequently investigated whether the ICD is sufficient to induce BDNF-mediated neurotrophic effects independently of the SorCS2-ECD. To isolate the SorCS2-ICD effects from the ECD, we fused the transmembrane (TM) region and the ICD of SorCS2 to an EGFP construct rendering a membrane anchored wild-type ICD (Fig. 3d and S6a). The chimeric EGFP-TM-WT variant was able to rescue BDNF-induced neuronal branching in *Sorcs2* KO neurons, suggesting that the SorCS2-ICD can act in the BDNF pathway independently of its extracellular domain (Fig. 3e-f). As we had observed that serine-to-alanine substitutions in the SorCS2-ICD caused loss of neurotrophic signaling, we speculated whether converse phosphomimetic mutations of the triple-serine motif could potentiate BDNF signaling. To this end, we designed a phosphomimetic (PM) ICD using serine-to-aspartate substitutions that structurally mimic the presence of a phosphoserine [34, 35] (Fig. 3d and S6a). Indeed, in *Sorcs2* KO neurons, the phosphomimetic EGFP-TM-PM showed an increased response to BDNF compared to SorCS2 FL and EGFP-TM-WT (Fig. 3e-f and S6b), suggesting that phosphomimetic modulation of the SorCS2-ICD can potentiate neurotrophic effects of BDNF.

We next investigated whether an isolated non-membrane bound, cytosolic SorCS2-ICD could rescue BDNF-induced neuronal branching in the absence of SorCS2. To test this, we cloned plasmids encoding the WT and phosphomimetic SorCS2-ICDs (S2sol-WT and S2sol-PM) without transmembrane regions (Fig. 3g and S6a) and cotransfected them into *Sorcs2* KO neurons with GFP. Here, BDNF increased branching of *Sorcs2* KO transfected with S2sol-WT compared to GFP controls, indicating that the SorCS2-ICD does not need membrane anchorage to mediate neurotrophic effects. Interestingly, transfection with S2sol-PM significantly increased neuronal branching even in the absence of BDNF, suggesting that the cytosolic, phosphomimetic SorCS2-ICD itself is capable of eliciting neurotrophic signals (Fig. 3h-i).

Taken together, these data suggest that the triple serine motif of the SorCS2-ICD is a key component of the BDNF/TrkB signaling pathways and that phosphomimetic manipulation of this can activate dendritic branching even in the absence of BDNF.

### The soluble SorCS2-ICD facilitates subcellular trafficking of TrkB

Downstream signaling in neuronal cells involves retrograde trafficking of signaling endosomes containing receptor/ligand complexes (such as BDNF/TrkB) associated with adaptor and effector proteins [36–38]. To understand whether the soluble SorCS2-ICD might affect receptor tyrosine kinase trafficking, we evaluated the effect of S2sol-WT and S2sol-PM on TrkB subcellular localization in co-transfected HEK293 cells by confocal imaging. We observed a marked change in subcellular localization when co-transfecting with TrkB and S2sol-WT or S2sol-PM compared to an empty vector (EV), as TrkB changed from membrane-localized to a perinuclear localization (Fig. 4a). We next explored whether this was also the case with endogenous TrkB in hippocampal neurons. For this, we co-transfected *Sorcs2* KO neurons with GFP (for visualization) and either the S2sol-WT or the S2sol-PM variant (or EV as a control) and immunostained for endogenous TrkB. We observed a considerable increase in TrkB signal intensity when transfecting neurons with the S2sol-WT and the S2sol-PM variant (Fig. 4b) suggesting increased overall TrkB levels.

**Fig. 4:**
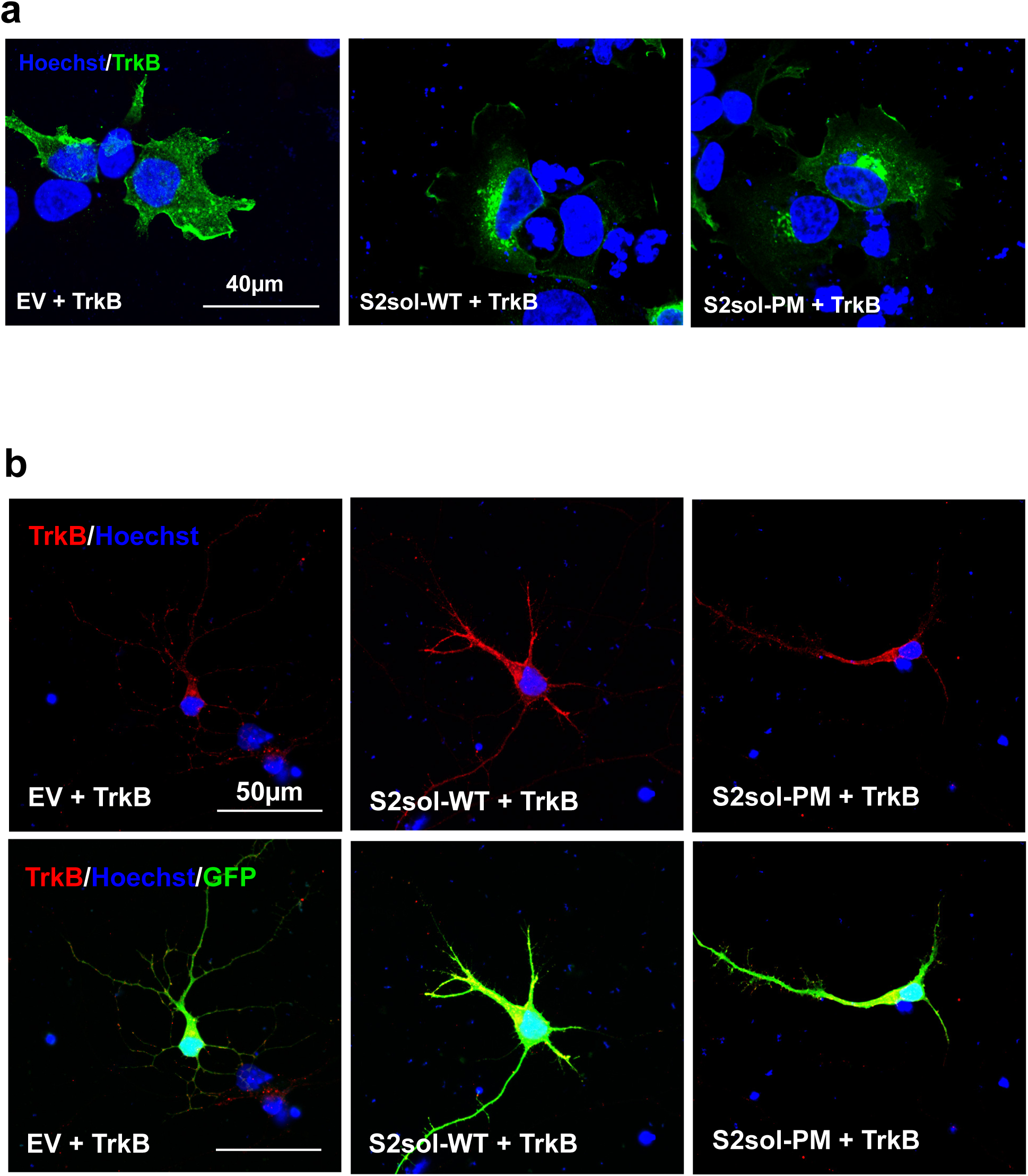
SorCS2-ICD regulates subcellular trafficking of TrkB. **(a)** Immunostaining of TrkB in HEK293 cells co-transfected with FL human TrkB together with empty vector (EV), S2sol-WT, or S2sol-PM, respectively. **(b)** Staining for endogenous mouse TrkB in *Sorcs2* KO hippocampal neurons co-transfected with GFP and soluble SorCS2 mutated (S2sol-PM) or wt tail (S2sol-WT) constructs. An empty vector was used in the control group.

### The neurotrophic effect of phosphomimetic activation of the ICD is conserved among SorCS1-3

Sequence alignment showed that the critical phosphoserine motif identified in the SorCS2-ICD is also present in SorCS1 and partly in SorCS3 (Fig. 2f). For this reason, we hypothesized that phosphomimetic modulation of the SorCS1 and SorCS3 ICDs could also induce neurotrophic activity. We therefore generated phosphomimetic, soluble ICDs for SorCS1 (S1sol-PM) and SorCS3 (S3sol-PM) (Fig. 5a and S6c) and similarly conducted dendritic branching experiments using these constructs in *Sorcs2* KO hippocampal neurons. Here, S3sol-PM increased neurite branching similarly to S2sol-PM, whereas S1sol-PM transfected *Sorcs2* KO neurons did not show significantly increased branching compared to the control (Fig. 5b-c, S6b).

**Fig. 5:**
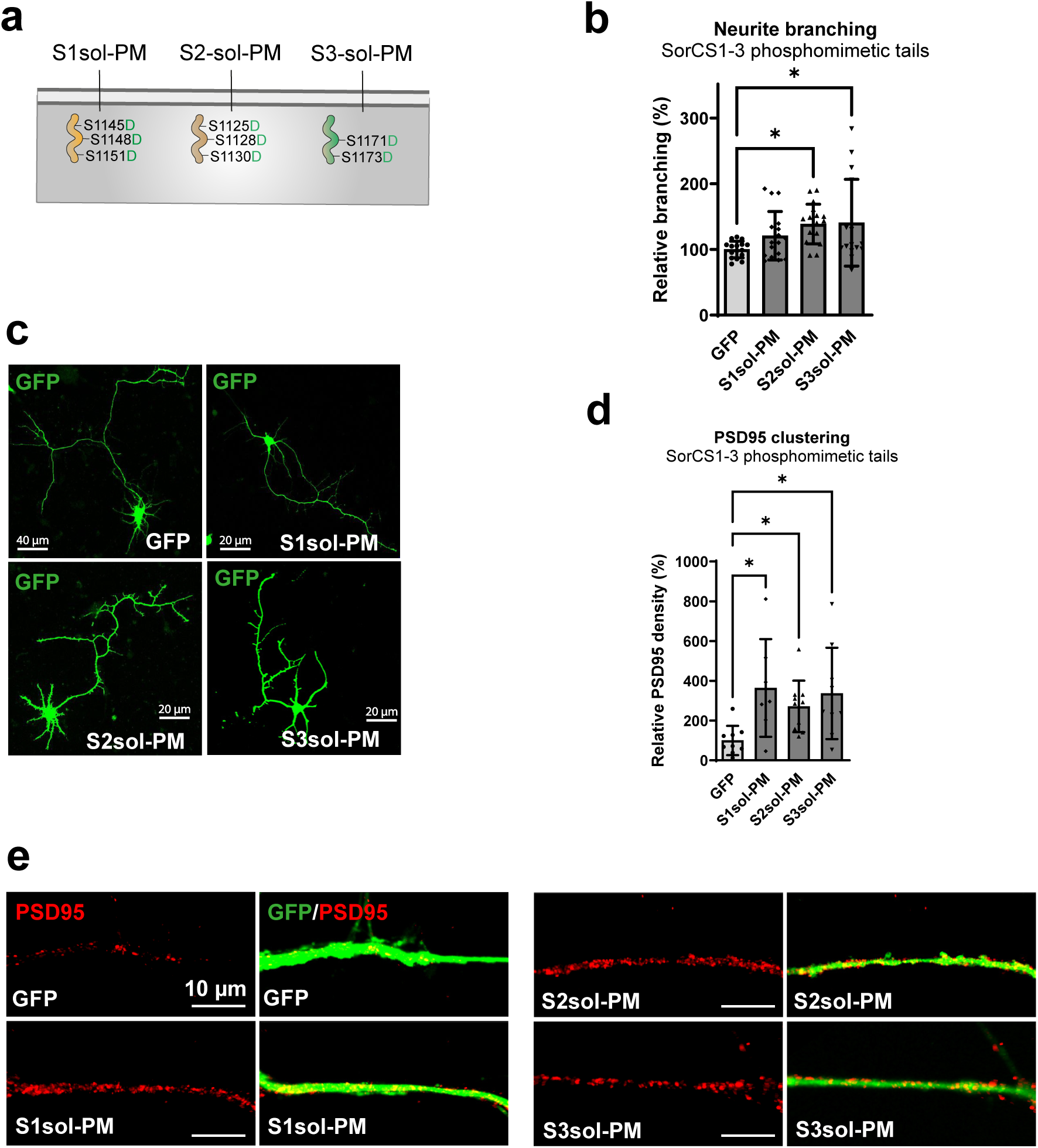
Soluble SorCS1-3 phospho-mimetic tails display neurotrophic activity. **(a)** Schematic of soluble SorCS1 (S1sol-PM), SorCS2 (S2sol-PM) and SorCS3 (S3sol-PM) tails with serine to aspartic acid substitutions as phospho-mimetic motif. **(b)** Neurite branching of *Sorcs2* KO hippocampal neurons transfected with soluble SorCS1-3 mutated tail construct (n=16-18 coverslips per group with 5-15 neurons analysed per coverslip) and **(c)** representative images. **(d)** PSD95 density measured in *Sorcs2* KO hippocampal neurons transfected with soluble SorCS1-3 mutated tail construct (n=6-8 coverslips per group with 15-20 neurites analysed per coverslip) and **(e)** representative images of PSD95 clusters in neurites. ROUT test was performed in PDS95 assay. Significance of neurite branching experiments and PSD95 density was calculated using ordinary one-way ANOVA (*p < 0.05). Data is represented as mean ± SD.

As SorCS1-3 regulate synaptic function and control correct synaptic protein assembly in response to neuronal activity [13, 18, 19, 21, 26], we further investigated the potential of the phosphomimetic soluble SorCS-ICDs to induce clustering of PSD-95 in *Sorcs2* KO hippocampal neurons as a measure of early synaptogenesis. Here, all constructs induced accumulation of PSD-95 in dendrites, suggesting they can initiate postsynaptic differentiation (Fig. 5d-e). Together, this suggests that the conserved triple serine motif in SorCS1-3 is involved in neurotrophic activities.

### Cell-penetrating, phosphomimetic triple serine motif peptides display neurotrophic activity

We speculated if the triple serine motif could be used as a basis for developing neurotrophic compounds. We therefore designed two phosphomimetic SorCS2-ICD-derived peptides: S2-LPep containing 8 amino acids from the C-terminal part of the transmembrane domain and the subsequent amino acids constituting TC2 (Fig. 2a); and S2-SPep encompassing the 21 amino acids that were deleted from TC2 to generate TC3. As a control, we developed a scrambled peptide from the S2-LPep construct (Scrambled). All peptides contained an N-terminal cell-penetrating peptide consisting of an 11 amino acid non-naturally occurring TAT-like transduction moiety to allow intracellular delivery [39] (Fig. 6a).

**Fig. 6:**
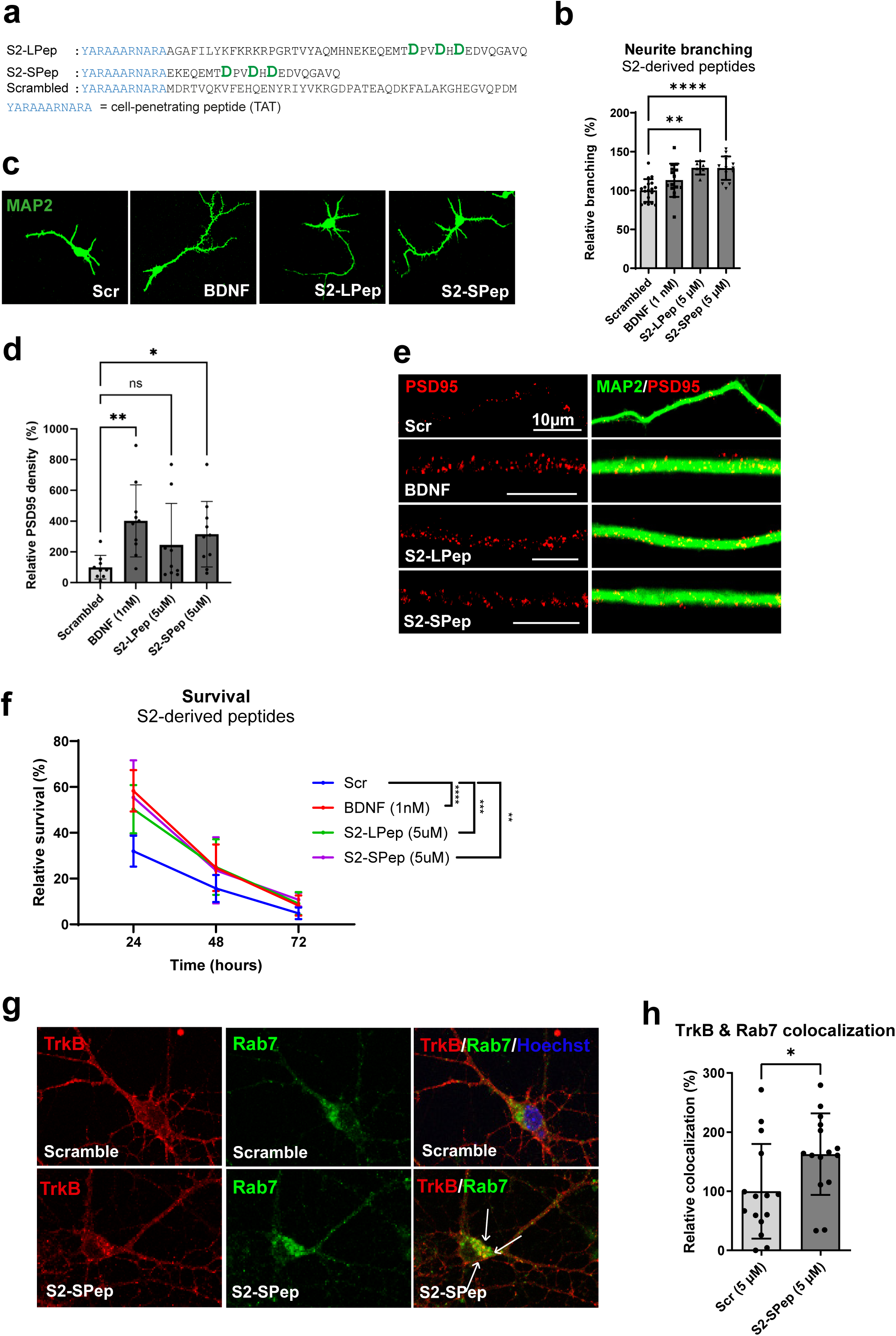
SorCS2 phospho-mimetic peptides induce neurotrophic response in neurons. **(a)** Sequences of SorCS2-derived peptides (short (SPep) and long peptide (LPep)) and scrambled peptide. Phospho-mimetic aspartic acids shown in green, TAT-sequence are marked in blue. **(b)** Neurite branching of wild-type (WT) hippocampal neurons treated with outlined compounds and **(c)** representative images (n=6-18 coverslips per group with 15 neurons analysed per coverslip). **(d)** PSD95 density measured in WT hippocampal neurons treated with outlined compounds (n=8-10 coverslips per group with 15-20 neurites analysed per coverslip) and **(e)** representative images of PSD95 clusters in neurites. **(f)** Relative survival of WT hippocampal neurons treated with outlined compounds over a period of 72 hours (n=4-8). **(g)** Representative confocal images of TrkB and Rab7 following S2-SPep or scrambled peptide treatment (5 µM, 10 min) and **(h)** the relative weighted colocalization calculated by the Zeiss Zen software (n=15). Significance of all experiments was calculated using two-tailed Student’s *t*-test (*p < 0.05, **p < 0.01, ***p < 0.001, and ****p < 0.0001). Data is represented as mean ± SD.

The peptide compounds were first tested for the ability to induce neuronal branching in WT hippocampal neurons. Both peptide variants (5 µM S2-LPep or 5 µM S2-SPep) led to an increase in neuronal complexity compared to Scrambled (5 µM) treated neurons (Fig. 6b-c). The neurotrophic effects of S2-SPep also translated to increased clustering of the postsynaptic marker PSD-95 in WT hippocampal neurons (Fig. 6d-e). We next assessed the effects of S2-LPep and S2-SPep to increase the viability of neurons in a viability assay with neurons seeded at low-density. Here, neurons quickly degenerate and perish as they lack neurotrophic input, which can be compensated for by exogenous neurotrophins [40]. Accordingly, the addition of S2-LPep and S2-SPep increased neuronal survival over a period of 72 hours (Fig. 6f).

As described above, we found that inactivation of the SorCS2-ICD by serine-to-alanine mutation results in a SorCS2 variant unable to transduce the response to BDNF. To understand whether this was the case for SorCS2-ICD peptide mimetics, we designed an inactive version of the short SorCS2-ICD derived peptide by substituting serines for alanines (denoted S2-SPepA, Fig. S7a) and tested its effect on inducing WT neurite branching in the presence or absence of BDNF. Intriguingly, S2-SPepA did not induce any morphological changes and even abolished the effects of BDNF (Fig. S7b-c), in line with observed effects in the experiments described above using the phospho-inactive FL-SorCS2 receptor (Fig. 3b-c).

As we observed an effect of S2sol-PM on both the retrograde transport and levels of TrkB, we further evaluated whether this also applied to SorCS2-derived peptides. For these studies, only S2-SPep was used, as this displayed similar or better neurotrophic effects when compared to S2-LPep (Fig. 6 and S7). Initially, we evaluated the levels of TrkB by western blotting in TrkB-transfected HEK293 cells following S2-SPep treatment (5 µM). Here, we observed an increase in TrkB levels 1-4 hours post treatment (Fig. S8a-b). Retrograde transport of signaling endosomes containing BDNF/TrkB is dependent on the small GTPase, Rab7 [41]. To establish whether the S2-SPep might affect the retrograde trafficking of TrkB, we analysed TrkB/Rab7 colocalization in WT neurons treated with S2-SPep (5 µM) at two timepoints (10 min and 1 hour) by confocal imaging. S2-SPep treatment increased colocalization of TrkB/Rab7 at 10 minutes post treatment (Fig. 6g-h), whereas no change was observed 1 hour post treatment (Fig. S9a-b). Following external growth-factor stimuli and signaling, receptors are either trafficked for degradation or recycled to the surface, for the latter requiring the GTPase Rab11 in the case of TrkB [42, 43]. As no difference in TrkB/Rab7 colocalization was observed 1 hour post treatment with S2-SPep we evaluated TrkB/Rab11 colocalization at this timepoint, to determine if TrkB might be recycled to the plasma membrane. Indeed, S2-SPep treatment increased Rab11/TrkB colocalization (Fig. S9c-d), indicating that S2-SPep induces recycling of TrkB, thereby potentially circumventing degradative pathways in line with the observed increase of TrkB in HEK293 cells following S2-SPep treatment. These results highlight SorCS2-derived peptides as potential modulators of retrograde and recycling processes of signaling receptor-complexes to induce neurotrophic support.

### S2-SPep activates signaling cascades downstream of TrkB

To investigate potential mechanisms for the neurotrophic effects elicited by the phosphomimetic peptides, we treated hippocampal neurons with either S2-SPep, S2-SPepA or Scrambled peptides (10 min, 5 µM) and analyzed the neuronal lysates using the KINEXUS assay previously described. S2-SPep regulated 48 phospho-sites at a nominal significant cutoff at α= 0.1 of which 27 phospho-sites remained significant following FDR correction while S2-SPepA only regulated 15 phospho-sites at a nominal significant cutoff (p<0.1) of which 7 phospho-sites remained significant following FDR correction (Fig. S10a). S2-SPep decreased phosphorylation of several proteins within the BDNF-pathway including TrkB (Y516), RSK (S221/S227), Cofilin-1 (S3) and PRKACA (T196/198) as well as several other kinases such as GSKa (T19/S21), MET (Y1234/Y1235/S1236), and CDK1 (T14) and increased phosphorylation of well-established BDNF-pathway proteins Src (Y419) and NMDAR1 (S896) as well as SIK3 (T163), Cyclin E1 (T395) and the protein tyrosine kinase selective ubiquitin ligases Cbl (Fig. 7a). Only MERTK (Y749) was regulated by both S2-SPep and S2-SPepA (Fig. S10b). KEGG pathway analysis of the nominally significant S2-SPep phospho-targets showed engagement in ERBB, MAPK and neurotrophin signaling among the most enriched pathways (Fig. 7b). STRING analysis of FDR significant hits further identified Src and TrkB as key components of the signaling events elicited by the S2-SPep (Fig. 7c).

**Fig. 7:**
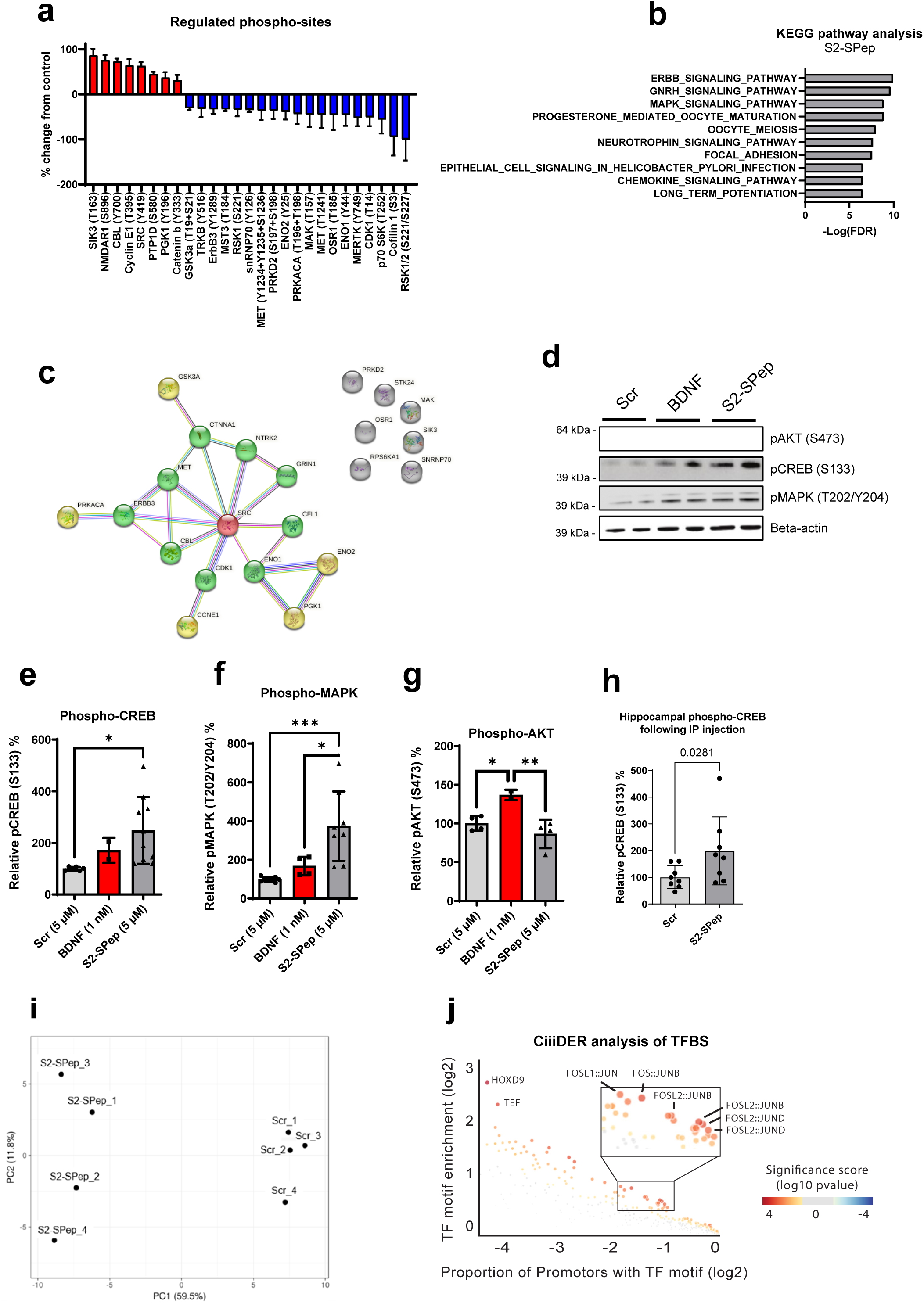
S2-SPep induces CREB activity both *in vitro* and *in vivo*. **(a)** Identification of kinase targets of S2-SPep (5 µM) relative to Scrambled-treated (5 µM) wild-type (WT) hippocampal neurons (DIV5) using a phospho-antibody array (n=3). Only significant (p<0.05) targets with ±30% change from control and signal intensities above 100 are shown. **(b)** KEGG pathway enrichment analysis of phospho-targets identified. **(c)** STRING analysis using high-confidence score (0.7) of identified phospho-targets **(d)** Western blots of phospho-CREB, phospho-MAPK, phospho-AKT and beta-actin in hippocampal neurons stimulated with Scrambled peptide (5 µM), BDNF (1 nM) or S2-SPep (5 µM) for 10 min and **(e-g)** densitometric analysis of phospho-proteins shown in **(d)** relative to beta-actin (n=2-4 for BDNF, n=4-8 for S2-SPep and Scrambled). Experiment was carried out twice in 2-4 replicates. Significance was calculated using two-tailed Student’s *t*-test (*p < 0.05, **p < 0.01 and ***p < 0.001). **(h)** WT mice injected with S2-SPep (∼36 mg/kg) intraperitoneally and subsequent (40 min. post injection) measure of phospho-CREB in hippocampal tissue by western blotting. Experiments were carried out three times with 2-3 mice per experiment per group (n=8). Significance was calculated using non-parametric Mann-Whitney U test. **(i)** Principal component analysis of mRNA sequencing data from wild-type hippocampal neurons treated with S2-SPep (5uM, 2 hours) or Scrambled peptide (5uM, 2 hours) (n=4). **(j)** Enrichment analysis of transcription factor binding sites in nominally significant genes using CiiiDER. Data is represented as mean ± SD.

S2-SPep activation of the MAPK pathway was further validated in WT neurons by western blotting. Here, S2-SPep activated both MAPK (T202/Y204) and its downstream target CREB (S133) (Fig. 7d-f). In line with the pathway analysis, S2-SPep did not activate Akt measured by S473 phosphorylation (Fig. 7g). Finally, we aimed to investigate whether the S2-SPep could engage these signaling pathways *in vivo*. We administered the S2-SPep or Scrambled peptide by intraperitoneal injection in WT mice at 36 mg/kg. Subsequent western blot assessment of hippocampal lysates showed a significant increase in phosphorylation of S133 CREB indicating that the SorCS2-ICD peptide mechanism is functional *in vivo* (Fig. 7h).

To further investigate the molecular consequences of S2-SPep induced signaling we studied the transcriptional changes elicited in neurons. WT neurons were stimulated with either S2-SPep or scrambled peptide (5 µM) for 2 hours and total RNA was subsequently isolated and sequenced. A clear effect of S2-SPep treatment on overall gene expression was apparent, with a Principal Component Analysis plot distinctly differentiating between the two treatments (Fig. f7i). More than 100 genes were differentially expressed in neurons exposed to S2-SPep compared to Scrambled at a nominal significant cutoff at alpha = 0.005 (Fig. S11b) of which 28 genes remained significant following FDR correction (Fig. S11a). Remarkably, transcription factor binding site (TFBS) analysis of the nominally significant genes revealed a strong enrichment of consensus motifs for Jun:Fos TFBS, which comprises the Activator Protein 1, AP-1. In addition to this, motifs for developmental transcription factors HOXD9 and TEF were also significantly enriched (Fig. 7j). Gene Ontology analysis of the nominally significant genes showed enrichment in signaling mechanisms including neurotransmitter release and response to cAMP, the key regulator of CREB activity (Fig. S11d). Further pathway analysis using KEGG showed that S2-SPep acts on gene networks involved with neurodegenerative disorders such as Parkinson’s Disease, Huntington’s Disease and Alzheimer’s Disease. Additionally, the engaged pathways involved mitochondrial energy processes of which BDNF/CREB are key regulators [44, 45](Fig. S11e).

## Discussion

The roles of the SorCS1-3 in CNS biology have been studied for many years, especially with the use of transgenic mice and primary neuronal cultures [46]. However, the molecular mechanism by which they exert their functions have not been elucidated. Here we propose that the activity of the SorCS1-3 receptors is governed by a conserved serine motif in their ICDs, which to our knowledge provides the first mechanistic insight into SorCS1-3 mediated signaling. These findings are summarized in a hypothetical model describing the role of SorCS2 in neurotrophic signaling (Fig. 9).

**Fig. 8:**
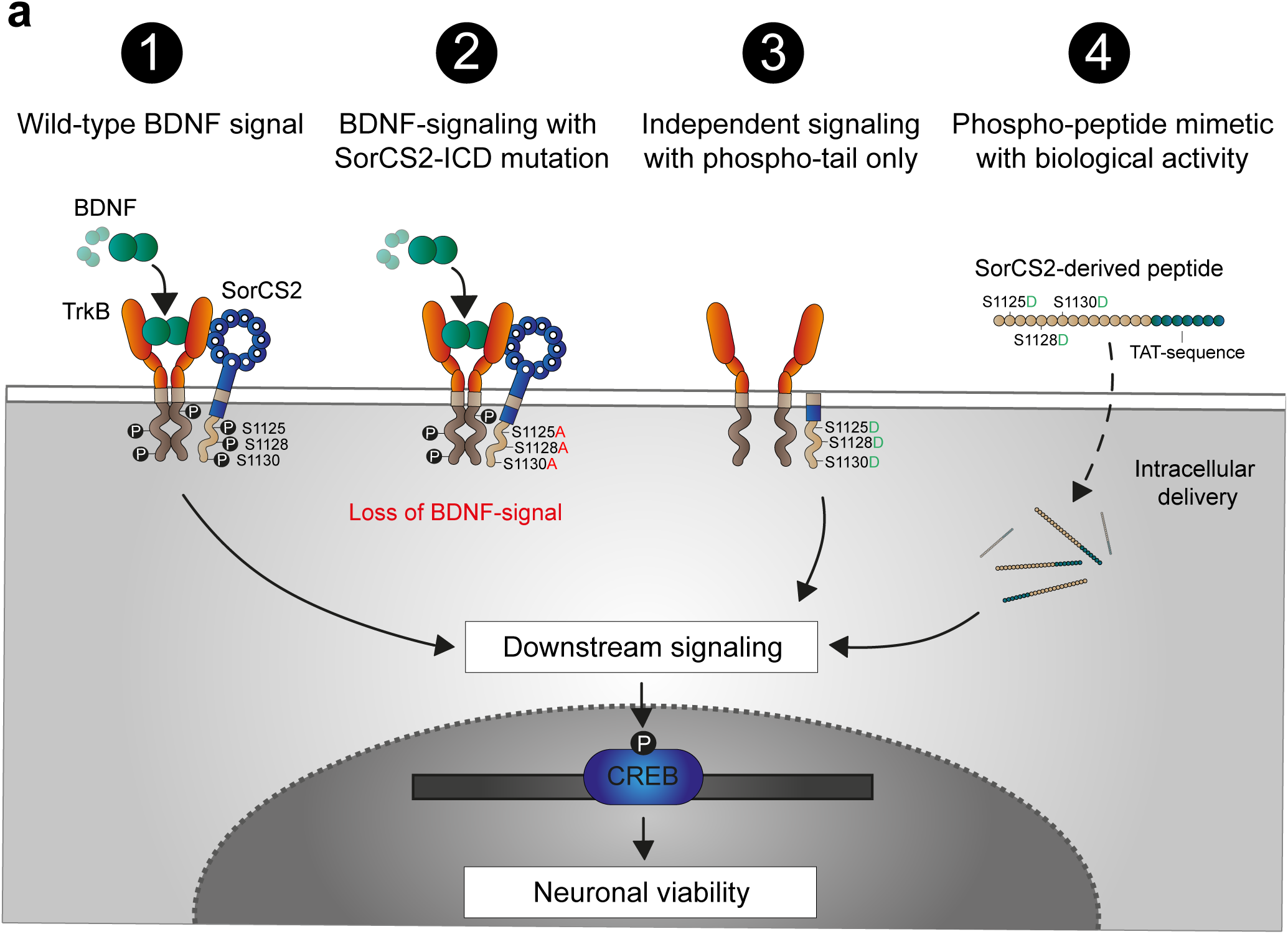
Hypothetical model describing the mechanistic basis for neurotrophic SorCS2-ICD peptide variants. **(a)** Schematic of the proposed model of SorCS2-ICD in BDNF-signaling in hippocampal neurons: **1)** In wild-type neurons, BDNF binds TrkB-SorCS2 complex at the post-synaptic site to induce phosphorylation of the intracellular domains of both TrkB and SorCS2 leading to downstream signaling and CREB activation. **2)** Mutation of SorCS2-ICD serine motif to alanine to elicit a dephosphorylated state leads to loss of BDNF signaling. **3)** A SorCS2 phospho-mimetic tail construct lacking the extracellular domain is capable of inducing a BDNF-like response even in the absence of BDNF. **4)** Phospho-mimetic peptides of the serine motif attached to a cell-penetrating moiety (TAT-sequence) elicit biological activity in neurons by activating CREB and induce neuronal viability independent of BDNF.

Previous studies have proposed that SorCS2 participates in pro and mature neurotrophin signaling by heterocomplex formation with either p75NTR or TrkB through its ECD. In this study, we find that loss of SorCS2 dramatically alters the phospho-kinase response to the neurotrophin BDNF in neurons, an effect that culminates in loss of neuronal branching. We identify a critical triple serine motif within the ICD of SorCS1-3, which is predicted to be phosphorylated, and indicate that SorCS1-3 could be targets of intracellular kinases as an immediate modulatory effect following TrkB activation. Extensive efforts were made to visualize SorCS2-ICD phosphoserines using trypsin-based nano LC-MS/MS, but unfortunately, we could only detect trace amounts of the unmodified target peptide most likely due to large peptide fragments rendered by trypsin digestion. Instead, we used ^32^P-labelling methods combined with SorCS2 immunoprecipitation, which indicated that SorCS2 indeed undergoes phosphorylation. Several other proteins that were co-precipitated with SorCS2 showed increased ^32^P signal, suggesting that adaptor proteins or other SorCS2 ligands might also be phosphorylated in response to BDNF. Furthermore, previous phosphoproteomic studies have demonstrated that SorCS1-3 indeed undergo phosphorylation on both intracellular and extracellular sites, but we have not found any reports of phosphorylations within the specific serine motif identified here. This could again be due to the methodological limitations of trypsin digestion and LCMS.

To substantiate our claim that the triple serine motif is indeed important for neurotrophic signaling, we initially generated serine-alanine variants of SorCS2. These variants generally lost their neurotrophic potential even though SorCS2 retained its extracellular domain, suggesting that this serine motif is critical for BDNF mediated effects. Notably, the S1125A SorCS2 mutant retained some activity following BDNF-stimulation, while no significant effects were observed for S1128A and S1130A mutants. As SorCS3 lacks the predicted phospho-serine at the corresponding S1125 position, it is our conjecture, that SorCS1-3 receptor activity is more likely regulated by the two latter serine residues of the motif for downstream target engagement. As the soluble, phosphomimetic serine-aspartate SorCS2-ICD variant showed neurotrophic effects independently of BDNF, we speculate that this triple serine motif possibly become phosphorylated mediating its neurotrophic effects.

Vps10p-domain receptors are known for their roles in subcellular trafficking of ligands, a function dependent on adaptor-binding motifs in their ICDs. Interestingly, we observed that a soluble SorCS2-ICD enables both the retrograde trafficking and the overall levels of TrkB. Modulation of intracellular serine motifs in other type I transmembrane receptors, has been shown to regulate the association with kinesin-1 and regulate trafficking [47] as well as proteolytic turnover [48–50]. Following an external stimulus, RTKs and their co-receptors are endocytosed to form a Rab7-positive signaling endosome, which either undergoes degradation or recycling to the surface [51, 52]. Interestingly, *Sorcs2* KO neurons display increased surface expression of Rab7 [18]. This led us to hypothesize that SorCS2 is involved in the formation and maturation of TrkB/Rab7-signaling endosomes, potentially dictated by the phosphorylation status of the SorCS2-ICD. In addition, this process might also trigger TrkB recycling rather than degradation, evident from our studies on overall TrkB levels. Indeed, this regulatory sorting mechanism through ICD modulation has been shown for the Vps10p-domain receptor SorLA on APP [53].

Our studies of phosphomimetic ICDs of SorCS1 and SorCS3 suggest that the observed function of the SorCS2 serine motif is preserved across the SorCS1-3 and that their activity is similarly governed by intracellular phosphorylation. However, the SorCS3-ICD contains a PDZ binding domain not found in SorCS1-2 [21] and the SorCS1- and SorCS2-ICDs undergo alternative splicing [23, 54] suggesting that heterogeneity at other sites in their ICDs might specify diverse interaction complexes of the receptors and their subcellular distribution, whereas the phosphorylation of the serine motif determines receptor activity.

Aberrant neurotrophic signaling is a hallmark of several neurological and psychiatric disorders. We therefore investigated if our findings could inform the development of neurotrophic active compounds by designing phosphomimetic peptides derived from the SorCS2-ICD. We observed effects on several parameters of neuronal health including neurite outgrowth, PSD-95 clustering at dendrites and neuronal survival. Interestingly, the SorCS2-ICD phosphomimetic peptides showed engagement with the MAPK-signaling pathway and induced CREB activation both *in vitro* and *in vivo*. Although the kinase array analysis showed strong engagement with BDNF-related pathways, the functional role of individual phosphorylation sites is not easily interpreted. As the assay was limited to a singular timepoint, a more detailed and time-resolved investigation might elucidate the dynamics of kinase signaling affected by the S2-SPep.

RNA sequencing further indicated that the identified DEGs were highly enriched in transcription factor binding sites for c-Fos:Jun heterodimers, in line with increased CREB activation leading to the expression of CRE-containing genes [55–57]. Both CREB and Fos:Jun heterodimers directly bind to mtDNA to regulate the expression of mitochondrial genes in neurons, which in the case of S2-SPep resulted in a general upregulation of mitochondrial genes [45, 58, 59]. The activity of the AP-1 complex, formed by c-Fos:Jun heterodimers, is directly modulated by a stress response through active isoforms of JNK, which is one of the major signaling cassettes in the MAPK-pathway in addition to ERK. Intriguingly, ERK and JNK are inversely regulated and display opposite effects on neuronal survival [60]. A BLAST search of the wildtype SorCS2-sequence covered by the S2-SPep revealed an intriguing alignment with a region in the JNK-binding domain in the scaffold protein JIP2 (or MAPK8IP2) (Fig. S12). This fragment serves as a selective scaffold module which interlinks JNK components to form a functioning JNK signaling module [61]. Notably, JIPs are also involved in regulating the canonical BDNF/TrkB pathway by both bridging TrkB-kinesin interactions to regulate TrkB anterograde transport and further by binding to and modulating the activity of Src and NMDAR-mediated signal transduction [62–64]. Together, it could be hypothesized that S2-SPep, and in consequence the phosphorylated SorCS2-ICD, might regulate multiple functions, including Src and NMDAR modulation, TrkB trafficking and the formation of a JNK signaling complex, leading to activation of ERK and CREB and downstream modulation of the AP-1 transcriptional complex. Intriguingly, S2-SPep did not affect Akt phosphorylation status and therefore seems to only participate in a subset of the signaling pathways induced by BDNF/TrkB activation.

Taken together, the role of the triple serine motif represents the first mechanistic understanding of how SorCS receptors play a role in neuronal signaling cascades. We show that phosphomimetic manipulation of the ICD is sufficient to induce neurotrophic signaling without exogenous BDNF or the SorCS2-ECD. Accordingly, phosphomimetic peptides of the SorCS2-ICD, delivered intracellularly using a cell-penetrating peptide linker, activate neurotrophic signaling in neurons, suggesting that modulators of SorCS triple serine motif may be a therapeutic strategy in neurological and psychiatric disorders.

## Acknowledgements

The present study is supported by the Danish Council of Independent Research Sapere Aude starting grant (SG, grant number DFF 4183-00604), the Novo Nordisk Foundation (SM), the Lundbeck Foundation (SG and MK R263-2017-3678).

We thank Mette Singers and Mitra Shamshali for their excellent technical support. Also, we are grateful for assistance from the Bioinformatics Core Facility of Aarhus University Health.

## Author contributions

AD and MK wrote the manuscript. LMB and SP conducted and analysed *in vitro* protein phosphorylation experiments in response to BDNF. PSM designed and produced plasmid constructs. MK, SM and AD conducted and analysed morphology studies. AD carried out TrkB regulation experiments. AD conducted and analysed all peptide stimulation experiments. MK and AD performed *in vivo* experiments. AD and PQ performed RNA-sequencing analysis and bioinformatics. SG and SM supervised the study. All authors read and approved the final manuscript.

### Corresponding authors

Correspondence to Simon Glerup or Simon Mølgaard.

### Data availability

RNA sequencing data and all graph raw data are available for review.

## METHODS

### CONTACT FOR REAGENTS AND RESOURCE SHARING

Further information and requests for resources and reagents should be directed to and will be fulfilled by the lead contact Simon Glerup

#### Mouse model

C57BL/6j BomTac wild type (WT) (Taconic) mice and *Sorcs2* KO mice on the same genetic background [1] were used for these studies. All experiments were approved by the Danish Animal Experiments Inspectorate under the Ministry of Justice (Permit 2011/561-119) and carried out according to institutional and national guidelines. All animals were bred and housed at the Animal Facility at Aarhus University. Animals were housed in groups of up to five mice per plastic cage (42 × 25 × 15 cm) under pathogen-free conditions with a 12-h light/12-h dark schedule and fed standard chow (Altromin #1324) and water ad libitum. Cages were cleaned every week and supplied with bedding and nesting material, a wooden stick, and a metal tunnel.

#### Neuronal cultures

Postnatal day 0 (P0) WT and *Sorcs2* KO pups were euthanized by decapitation, brains removed and hippocampi dissected into ice-cold Leibovitz’s L-15 Medium (Life Technologies #11415049). The dissected hippocampi were dissociated for 30 min in 20 U/mL pre-activated papain (Bionordika, #WBT-LS003126). Hereafter, the tissue was washed in DMEM (Lonza, #BE12-604F/U1) containing 0.01 mg/mL DNase (Sigma, DN25) and 10% Fetal Bovine Serum (FBS) (Gibco, #10270-106) and triturated in DMEM containing 0.01 mg/mL DNase I and 10% FBS. Next, DMEM was removed and Neurobasal-A Medium (Gibco, #10888022), containing B-27 Supplement (Gibco, #17504044), 2 mM GlutaMAX (Gibco, #35050), 100 μg/mL Primocin (Invivogen, #ant-pm-2) and 20 μM floxuridine (Sigma, #F0503) + 20 μM uridine (Sigma, #U3750), was added to the cells to optimize neuronal growth conditions and remove mitotic cells. The cells were seeded on poly-D-lysine (Sigma #P7886) for coverslips or poly-L-lysine (Sigma #P1524) for plastic surfaces in addition to laminin (Invitrogen, 23017-015) at a desired density and left at 37°C and 5% CO_2_, with medium change every second day, before being fixed in ice cold 4% PFA after the desired days *in vitro (DIV)*. This procedure was used for all experiments using primary hippocampal neurons in culture.

#### Constructs

Full length SORCS2 (A variant) (GenBank OQ616758) was expressed by inserting the coding sequence into pcDNA3.1/zeo(-) (Invitrogen). Truncated variants of SORCS2A terminating at positions: K1101, N1117, and L1152 were established by insertion of PCR fragments. A truncated variant of SORCS2B (GenBank no. OQ616759) terminating at position 1152L was also created and used in the truncation experiments. For subsequent mutagenesis and chimeric experiments, the SORCS2A variant was used. Full length SORCS2A cytoplasmic domain mutations: S1125A/S1128A/S1130A and S1125D/S1128D/S1130D were created using a QuikChange Lightning kit (Agilent). Chimeric proteins including a signal peptide, EGFP and SORCS2A spanning from Q1173-S1159 (transmembrane and cytoplasmic domains) and the derived S1125D/S1128D/S1130D mutated variant were expressed using the pSecTagB vector (Invitrogen). The SORCS2A cytoplasmic domain spanning from A1094 to S1159 and the derived variant harboring the mutations S1125D/S1128D/S1130D were expressed using the pCpGfree-vitroNmcs vector (Invivogen), sequences were preceded by an AUG codon. The SORCS3 (GenBank NM_014978.3) cytoplasmic domain spanning from L1138 to V1222 and the derived variant harboring the mutations S1172D/S1174D were expressed using the pCpGfree-vitroNmcs vector (Invivogen), sequences were preceded by an AUG codon. The SORCS1 (NM_001206569.2) cytoplasmic domain spanning from L1114 to K1179 and the derived variant harboring the mutations S1145D/S1148D/S1150D were expressed using the pCpGfree-vitroNmcs vector (Invivogen), sequences were preceded by an AUG codon. The pEGFP-N1 (Clontech) plasmid was applied for co-expression in neurons. TrkB isoform C (GenBank NM_001018064.3) was expressed using pCpGfree-vitroNmcs vector. The empty vector used was pcDNA3.1/zeo(-) (Invitrogen).

#### Transfection of Sorcs2 KO neurons for imaging

*Sorcs2* KO neurons were isolated from p0 pups and seeded at 100.000 pr coverslip. Concerning branching experiments, 24 hours post seeding the neurons were transfected with either one of the different variants of FL-SorCS2 or tail-versions or an empty vector, along with GFP using Lipofectamine™ (ThermoFisher, #A12621). After 6 hours, the medium was changed to standard neurobasal A medium and the cultures were left at 37°C and 5% CO_2_ for 72 hours before being fixed in ice cold 4% PFA for 20 minutes and subsequent neurite branching imaging. When assessing postsynaptic differentiation or TrkB localization, the neurons were transfected at DIV10 with either SorCS1, SorCS2 or SorCS3-tail variants along with GFP using same method as described above. Neurons were fixed at DIV 13. PSD95 and TrkB were subsequently visualized by immunostaining as described above.

#### Stimulation of WT neurons for imaging

WT neurons were isolated from p0 pups and seeded at 10.000 pr coverslip for neuronal branching experiments or 50.000 for synaptogenesis. Concerning branching experiments, 24 hours post seeding the neurons were treated by changing half the media with either S2-S2Pep, S2-S2PepA, Scrambled or BDNF to yield a final concentration of 5 µM of peptides and 1 nM of BDNF. The neurons were fixed at DIV4. Concerning postsynaptic differentiation experiments, the neurons were treated at DIV10 similar to branching experiments and fixed at DIV13. Concerning TrkB/Rab7 or TrkB/Rab11 colocalization experiments, the neurons were treated at DIV7 by changing all the media with 5uM S2-S2Pep or 5uM Scrambled and fixed at either 10 or 60 minutes post treatment.

#### Immunostaining of cells on coverslips

The coverslips with the fixed cultured primary hippocampal neurons or HEK293 cells were briefly washed in D-PBS (Gibco, 14190-094). The coverslips were washed three times 5 min in D-PBS containing 0.1% Triton X-100 at RT to permeabilize the cells. Coverslips were washed once in D-PBS to wash away excessive Triton X-100 before blocking in 0.5 mL D- PBS containing 10% FBS for 30 min at RT. The coverslips were then incubated with primary antibodies (1:500 goat anti human/mouse TrkB, R&D Systems #AF1494; 1:1000 rabbit anti-mouse MAP2, Millipore #AB5622; 1:200 mouse anti-mouse PSD95, Sigma #P246; 1:100 rabbit anti-mouse Rab7, Cell Signaling Technology #9367; 1:100 rabbit anti-mouse Rab11, Cell Signaling Technology #5589), diluted in D-PBS containing 10% FBS and left overnight at 4°C in a humidified chamber. All antibodies used have been reviewed at the independent antibody validation database (www.pAbmAbs.com). The following morning, the coverslips were left in the humidified chamber at RT for 1 h before continuing the immunostaining process. The coverslips were washed three times 5 min in D-PBS containing 0.1% Triton X-100. Hereafter, the coverslips were incubated with secondary antibody (1:300 donkey anti-goat 568, or 1:300 donkey anti-mouse 568), diluted in D-PBS containing 10% FBS, in a humidified chamber at RT for 4 hrs. Subsequently, the cells were washed in D-PBS three times 5 min, with 5 µg/mL Hoechst nuclear stain, included in the last wash. The coverslips were mounted using Dako Fluorescence Mounting Media, sealed with nail polish and stored at 4°C. This method was used for all immunostainings of cultured hippocampal neurons.

#### Confocal microscopy and image analysis

The immunostaining samples were analyzed on a Zeiss confocal LSM 780 or 510 microscopes using 20X/0.8 M27 and 63X/1.20 W Korr (Water immersion correction ring) objectives. Appropriate filters were used upon excitation of the different fluorophores to match their maximum fluorescence emission. The wavelengths used were 258, 488 and 568 nm and they were configured to obtain the best signal during image acquisition of the samples in order to prevent bleed through between the different probes. Confocal microscopy images were processed and performed in Zen 2011 Image Processing software (Carl Zeiss) or by Imaris software (Bitplane). All images compared were subjected to similar brightness and contrast adjustments. Rab7/TrkB and Rab11/TrkB colocalization was performed using integrated Weighted Colocalization Coefficients in the Zen software.

#### Neuronal branching assay analysis

Neuronal branching experiments were performed in two different systems to ascertain the effects of either overexpressed SorCS2 constructs or SorCS2 peptide mimetics.

##### Transfection

Due to the density of the culture as well as the low transfection efficiency with lipofectamine in postnatal neurons, we used GFP as a co-transfectant in all conditions to visualize neuronal morphology only of transfected neurons. As such, GFP co-transfected with an empty vector was used as a negative control. Branching was counted pr “branch order”: all dendrites extending from the soma were considered branch order 1, and this pertained to the entirety of the longest neurite extension for such 1^st^ order branches. Thus, any additional branches on a primary branch where considered a 2^nd^ order branch etc. As there are rarely any 5^th^ order branches in this assay, any such were annotated as 4^th^ order. The sum of all branches (independent of branch order) was used for the total number of dendritic branches.

##### Peptide assessment

Here, low density neuronal cultures (10.000 pr coverslip) were used to be able to distinguish projections from individual neurons. MAP2 staining was used to visualize dendrites. Neurons that were in close proximity were discarded from the analysis to exclude bias caused by neuron-neuron neurotrophic support. Branching was counted as for transfected neurons. Up to 25 neurons were imaged per coverslip and the mean number of branches was taken per coverslip. Coverslips with less than 5 transfected neurons were disregarded in the analysis.

#### Assessment of phospho-regulated proteins

Hippocampal neurons from p0 WT and *Sorcs2* KO pups were seeded at a density of 2 mio pr well as described earlier. After 7 DIV, the neurons were stimulated for indicated timepoints (10 minutes for phospho-array) with NBA media (no additives) containing either 1 nM BDNF or similar volume of sterile PBS. In relation to peptide-assays either 5uM S2-SPep, 5uM S2-SPepA or 5uM Scrambled was used. Hereafter, the neurons were washed once in ice cold PBS followed by lysis in ice cold KINEXUS special lyse-buffer (20mM MOPS, 2mM EGTA, 5mM EDTA, 50mM Sodium fluoride, 60 mM b-glycerophosphate (pH7.2), 25 mM sodium pyrophosphate, 2.5 mM sodium orthovanadate, 50nM phenylarsine oxide, 1% triton X-100 and 0.05% sodium dodecylsulphate (SDS)). 1mM DTT was added to the lysis buffer prior to use, and the samples were hereafter kept on ice. Samples where then sonicated four times for ten seconds and centrifuged followed by removal of cell pellet. BCA protein assay were performed to measure determine protein concentration and to ensure the samples met KINEXUS-demands for protein-concentration, and the samples were frozen and sent to KINEXUS for analysis or analysed by western blotting. The KAM-1325 antibody-microarray was used to screen for kinase-activity and comparisons were made between stimulated and un-stimulated wells within genotype.

#### SorCS2 phosphorylation

WT hippocampal neurons were seeded at a density of ∼2M per well according to previously described method. At 7 DIV the neurons were washed twice in DMEM without phosphate and incubated with 40 µCi/mL phosphorus-32 radionuclide (PerkinElmer, Specific Activity: 8500-9120Ci/mMole) in DMEM without phosphate for 4 hours at 37°C. After incubation, the cells were incubated with fresh 0.4 mCi/mL 32P DMEM with or without 1nM BDNF for another 10 minutes at 37°C and thereafter lysed on ice using KINEXUS lysis buffer (described previously). Lysates were spun down at 14000 rpm at 4°C for 10 minutes and supernatant were incubated with GammaBind Sepharose (GE Healthcare, # 17088501) beads premixed with 1µg/mL rabbit α-mouse SorCS2 F7378 (DAKO custom-made polyclonal ab against the cytoplasmic tail of mouse SorCS2) antibody overnight. Next day the beads were washed in PBS 3 times 5 minutes with 0.05% Tween-20 prior to boiling and denaturation by LDS and DTT (2:1 mixture) at 95°C for 10 min. The eluate was run by SDS-PAGE as previously described and the radiation was detected and analysed using fuji imaging plates and Image Reader FLA-3000 (FujiFilm).

#### SDS-PAGE and Western blotting

Cell lysates were spun down at 14000 rpm at 4°C for 10 minutes and cell pellets were discarded. Protein concentration of lysates was measured in triplicates using bicinchoninic acid (BCA) assay. 50 µg of protein was mixed with ¼ of total volume LDS Sample Buffer (Invitrogen, #NP0007) and 1/10 of total volume DTT. The protein mixture was boiled at 95 °C for 5 minutes before run on a 4-12 % Bis-Tris gel (Invitrogen, #NP0335BOX) in NuPAGE MOPS buffer (ThermoFisher, #NP0001) or NuPAGE MES buffer (ThermoFisher, #NP0001).

The proteins were transferred to nitrocellulose membrane using iBlot Transfer Stack kit (Invitrogen, NB301001) and iBlot Dry Blot System (Invitrogen, #IB1001). The membrane was blocked in Blocking buffer (Tris-base 50mM, NaCl 500mM, 2% Milk powder and 1% Tween-20) for at least 30 minutes and incubated with primary antibody (1:1000 rabbit anti-human/mouse phospho-CREB (S133), Cell Signaling Technology #9198, 1:1000 rabbit anti-human/mouse phospho-Akt (S473), Cell Signaling Technology #4060, 1:1000 rabbit anti-human/mouse phospho-MAPK (T202/Y204), Cell Signaling Technology #9101 and 1:5000 mouse anti-mouse beta-actin, Sigma #A5441) overnight (o/n). The antibodies were diluted in Blocking buffer. The next day, the membrane was washed three times for 10 minutes in washing buffer (CaCl2 2mM, MgCl2 1mM, HEPES 10mM, NaCl 140mM, 0.2% Milk powder and 0.5% Tween-20) and incubated with HRP-conjugated secondary antibody (diluted 1:2000 in Blocking buffer, rabbit α-mouse DAKO #0260, swine α-rabbit DAKO # P0217) for 1 hour at room temperature. Next, the membrane was washed 3x10 min. in washing buffer and bands developed using Amersham ECL Western Blotting Detection Reagent (GE Healthcare, # RPN2106) - following supplier’s instructions - and detected and analyzed using LAS-4000 and appurtenant software (GE Healthcare). The intensity of bands was quantified by densitometric analysis using Multi Gauge V3.2 software.

#### TrkB uptake assay

HEK293 cells were seeded at a density of 20.000 cells per coverslip. The cells were co-transfected the following day with soluble SorCS2 ICD-variants (S2sol WT, S2sol PM) and TrkB and incubated o/n. Next, the cell media were exchanged with 400 µL ice-cold DMEM media containing 10 % FBS and 1 µg/mL α-TrkB antibody (goat anti human/mouse TrkB, R&D Systems #AF1494). The cells were incubated on ice for 2 hours. Next, the cells were washed once in ice-cold DMEM with 10 % FBS. Medium was then exchanged with warm DMEM + 10 % FBS starting with cells that incubated at timepoint 60 min. then 40 min. and so on. At timepoint 0, the cells were put on ice, then fixed and stained for TrkB.

#### Survival assay

WT hippocampal neurons were seeded at a density of 2000 cells in a 35x12mm Dish (ThermoFisher #171099) with a 1 cm^2^ marked underneath. The neurons were allowed to settle for 3 hours and then counted in a light microscope within the marked area (giving total amount of neurons). Next, the cells were treated with either SorCS2-ICD derived peptides (1uM) or BDNF (1nM) by exchanging half the media with NBA containing B27, F+U and primocin. Living neurons were counted at 24 hours, 48 hours and 72 hours post seeding using a light microscope. Neurons were considered alive if having neuronal processes. The survival rate of the neurons was calculated as the number of neurons alive at the different timepoints divided by the total amount counted after seeding the cells.

#### In vivo experiments

The WT mice were received from Taconic. 36 mg/kg of S2-SPep was injected intraperitoneally (I.P). and 40 min. after injections, the mice were sacrificed by cervical dislocation and hippocampus tissue was taken and put in 250 µL of KINEXUS buffer on ice. The tissue was immediately lysed using TissueLyser II (Qiagen, #85300). This was followed by pelleting membranes and un-lysed tissue by centrifugation at 16000 rpm, 4 °C for 10 min. The pellets were discarded. 5 µL of 0.5 mM TCEP was finally added to the lysate and lysates were stored at −20°C.

#### RNA-seq analysis

Total RNA was isolated from hippocampal neurons using the RNeasy Mini Kit (Qiagen) following the manufacturer’s instructions. The quantity and quality of RNA was determined using an Agilent 2100 Bioanalyzer (Agilent technologies, SantaClara, USA). 8 RNA samples (200 ng RNA/sample; 4 samples/study group, RNA Integrity Number (RIN) > 7.30 (mean = 8.38, SD = 0.59, no significant difference between study groups) from primary hippocampal neuron culture were included for RNA-seq analysis. cDNA was synthesized from each individual sample using random hexamer primers and libraries were prepared by TruSeq RNA sample preparation kit (Illumina, San Diego, USA). RNA-seq analysis (50 bp; single-end; minimum 20 million clean reads/sample) was performed using BGISEQ-500 (BGI Global Genomics Services, Copenhagen, Denmark).

Single-end reads with more than 90% bases having less than 1% sequencing error and no ambiguous bases were aligned to the mouse genome assembly GRCm38.72 using HISAT2 v2.1.0 [2]. Overall rates of alignment were above 97% for all libraries. StringTie v1.3.4 [3] was applied to assemble transcripts and generate counts of fragments uniquely mapped to known and annotated genes using the Ensembl annotation file Mus_musculus.GRCm38.72.gtf. The count table of uniquely mapped fragments was used for differential expression analysis using DESeq2, Bioconductor v3.4.1 [4] using default parameters.

#### CiiiDER analysis

Enrichment analysis of TFBSs was carried out according to Gearing et al using CiiiDER [5]. Briefly, promotor sequences (2000 bp upstream of TSS) were extracted from the Homo sapiens GRCh38 – Ensemble release 108) genome file. Identification of TFBSs in these sequences was performed with JASPAR2020_CORE_vertebrates transcription factor position frequency matrices (downloaded from https://jaspar.genereg.net/) and a deficit cut-off of 0.15. CiiiDER enrichment analysis of overrepresented NR TFBSs in DEG query sequences compared to non-DEG query sequences (from 5000 genes with p∼1 and logFC∼0) was determined by comparing the number of sequences with predicted TFBSs to the number of those without, using a Fisher’s exact test.

#### Statistical analyses

Data are presented as mean ± SD. Statistical significance was evaluated using two-way ANOVA followed by post hoc analysis using Student’s t-test unless mentioned otherwise.

Adjusted p-values from RNA-sequencing analysis were calculated by Benjamini-Hochberg false discovery correction (5 %) for DEseq2. Genes with adjusted p-values less than 0.05 were considered as DEGs. Principal Component Analyses (PCAs) and cluster analyses were performed using Clustvis [6]. Functional analyses of DEGs were performed using Ingenuity Pathway Analysis (QIAGEN Bioinformatics, Redwood City, CA, USA).

Significant hits in KINEXUS data were calculated based on Change from Control (CFC), with adjustment for Error Range (ER). Hits for BDNF-induced phospho-hits in *Sorcs2* KO vs. WT were defined as CFC ≥ 45; SUM of ER <0.75 of CFC value; At least one Globally Normalized intensity value ≥ 1500. Hits for S2-SPep or S2-SPepA phospho-hits in WT neurons were defined as CFC ≥ 30; SUM of ER <0.75 of CFC value; At least one Globally Normalized intensity value ≥100.

STRING analysis was performed on significant phospho-site hits from the KINEXUS data described above. For the analysis, a high-confidence score of 0.7 was used.

Pathway enrichment (KEGG) & GO enrichment were carried out on nominal significant DEGs (p<0.005) and phospho-sites (p<0.1).

---

**Supplementary Figure 1:**
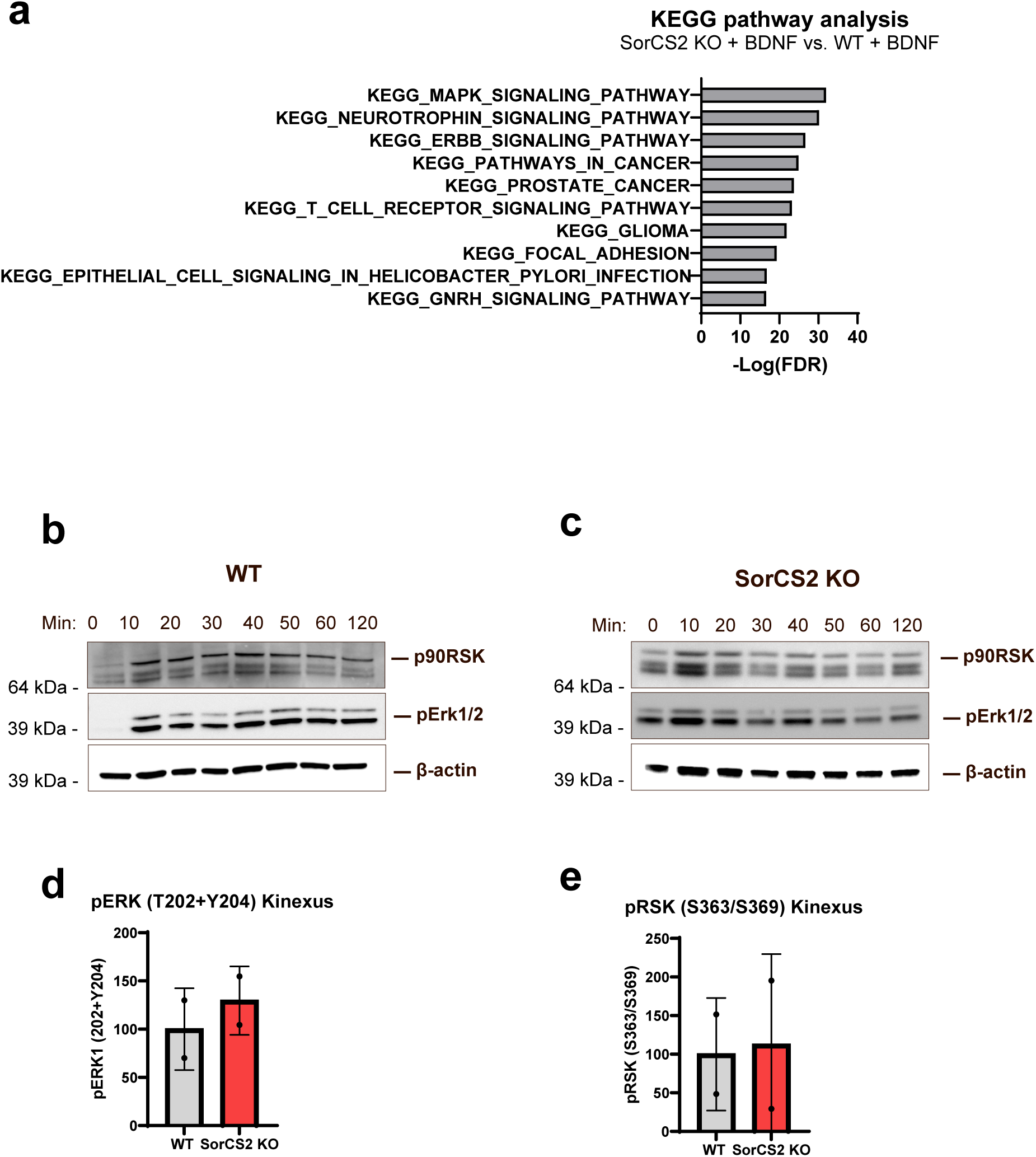
**(a)** KEGG pathway analyses of proteins significantly regulated between SorCS2 KO stimulated with BDNF vs. WT stimulated with BDNF. **(b-c)** Time course of BDNF-induced activation of MAPK and RSK in WT **(b)** and KO **(c)** hippocampal neurons analyzed by western blotting. WT and KO samples were run on separate blots and thus not directly comparable. **(d-e)** Normalized Kinexus signals for pERK **(d)** and pRSK **(e)** indicate no baseline differences between WT and *Sorcs2* KO. Data represented as mean + SD.

**Supplementary Figure 2:**
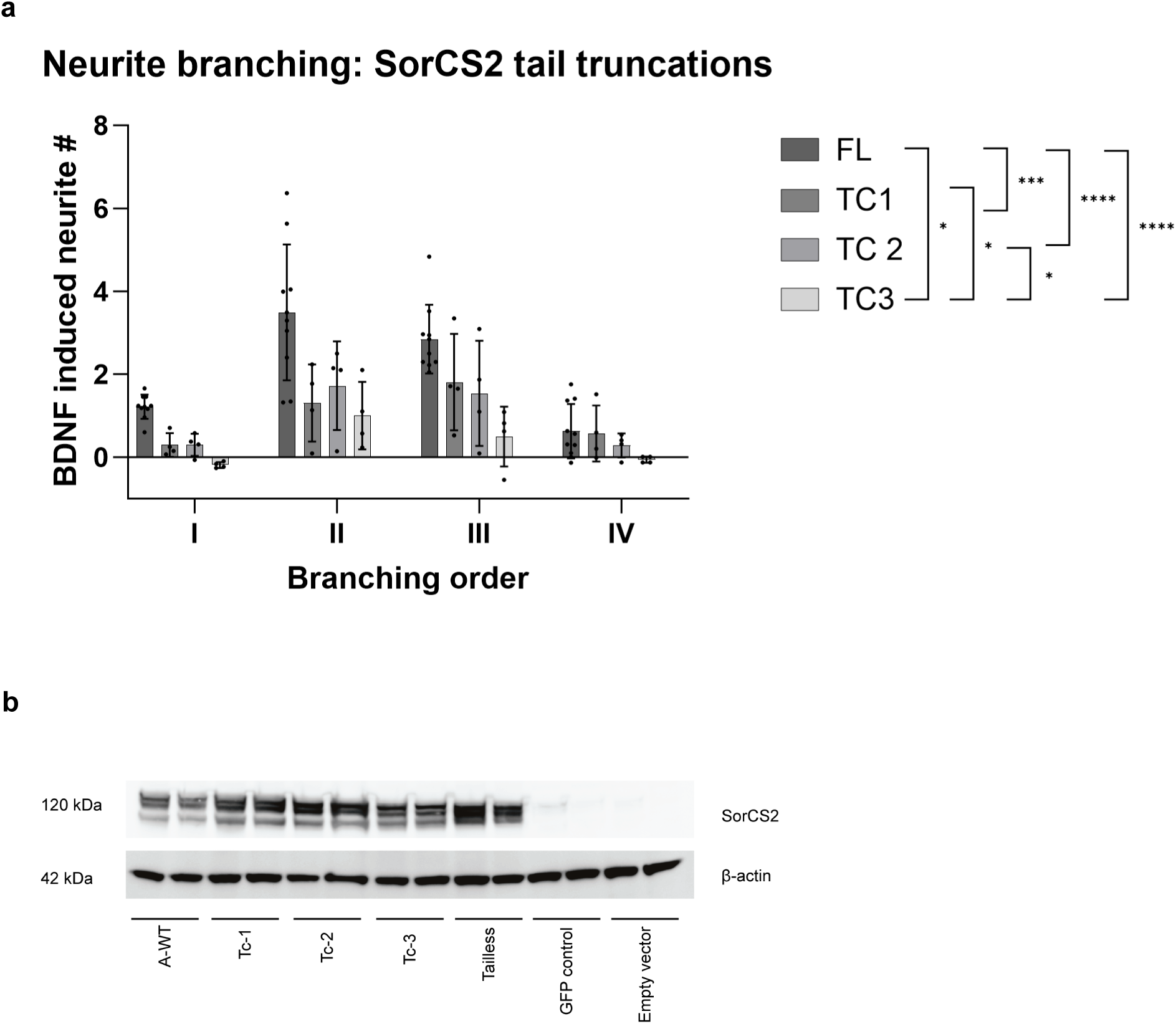
**(a)** Extended data on neuronal branching orders in response to BDNF (1nM) in *Sorcs2* KO hippocampal neurons co-transfected with GFP and truncated variants of SorCS2 receptor (n=4-6 coverslips per group with 5-15 neurons analysed per coverslip). Significance was calculated using ordinary two-way ANOVA of main effects. *p < 0.05, ***p=0.0001 and ****p < 0.0001. Data represented as mean ± SD. **(b)** Western blotting of lysates from HEK293 cells transfected with the various SorCS2 TC variants.

**Supplementary Figure 3:**
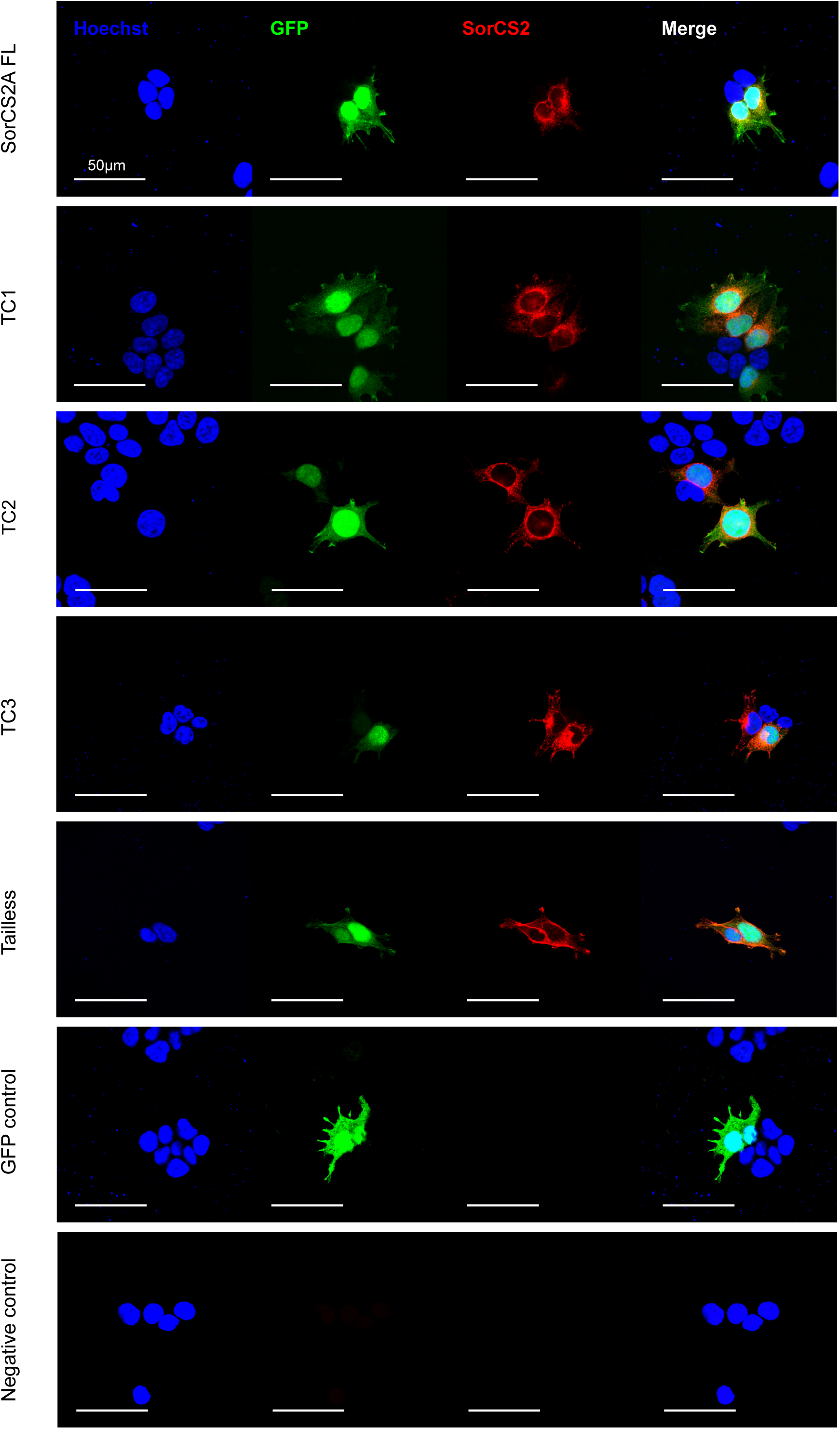
Immunocytochemistry stainings of SorCS2 in transfected HEK293 cells with GFP and truncated variants of SorCS2 receptor.

**Supplementary Figure 4:**
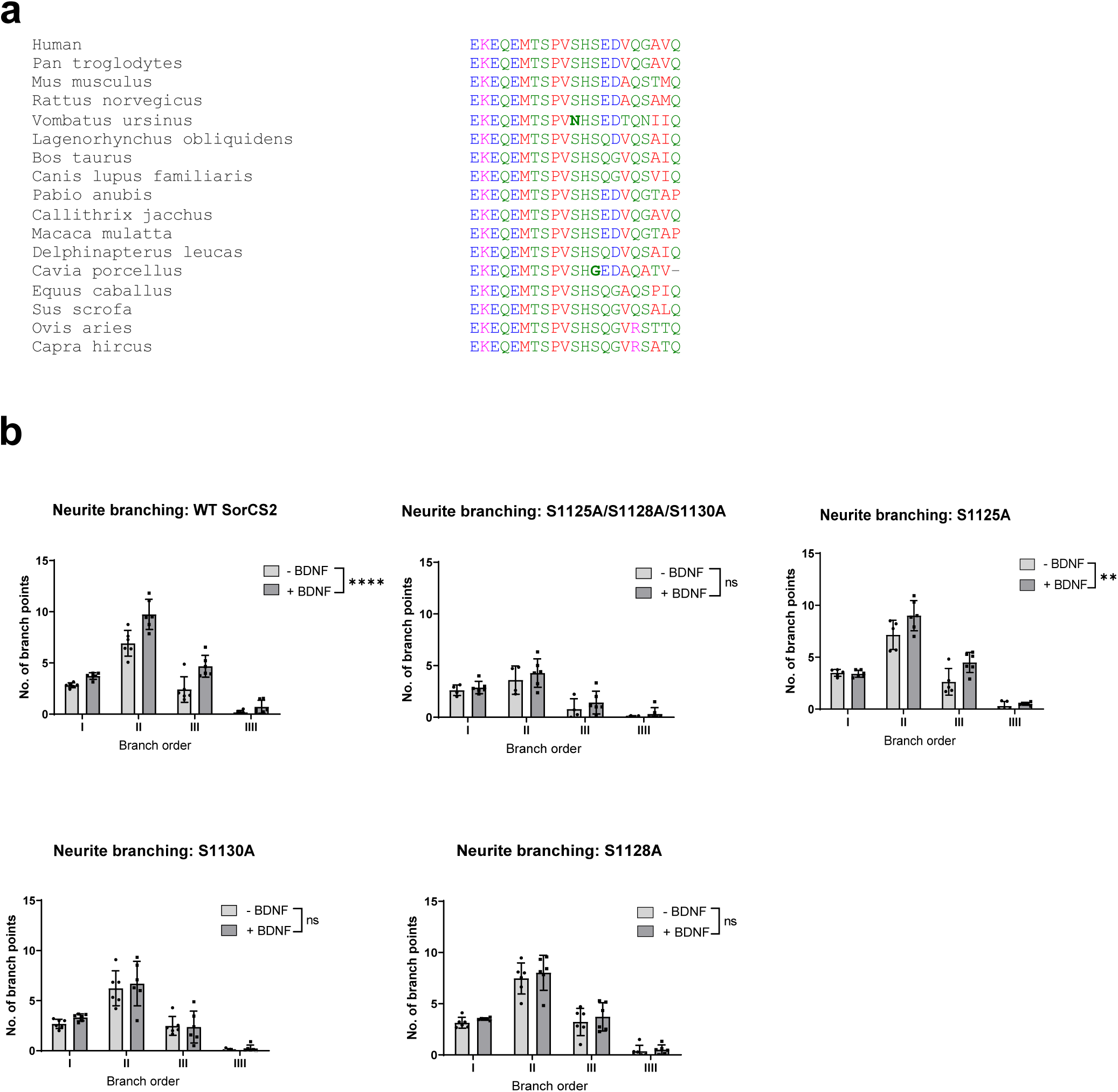
**(a)** Alignment of SorCS2-ICD region containing serine-motif across species. **(b)** Extended data on neuronal branching orders in response to BDNF (1nM) in SorCS2 KO hippocampal neurons co-transfected with GFP and SorCS2 variants with serine-to-alanine mutations (S1125A, S1128A, S1130A, S1125A/S1128A/S1130A) or wild-type SorCS2 (WT) (n=4-6 coverslips per group with 5-15 neurons analysed per coverslip). Significance was calculated using ordinary two-way ANOVA of main effects. **p < 0.01 and ****p < 0.0001. Data represented as mean ± SD.

**Supplementary Figure 5:**
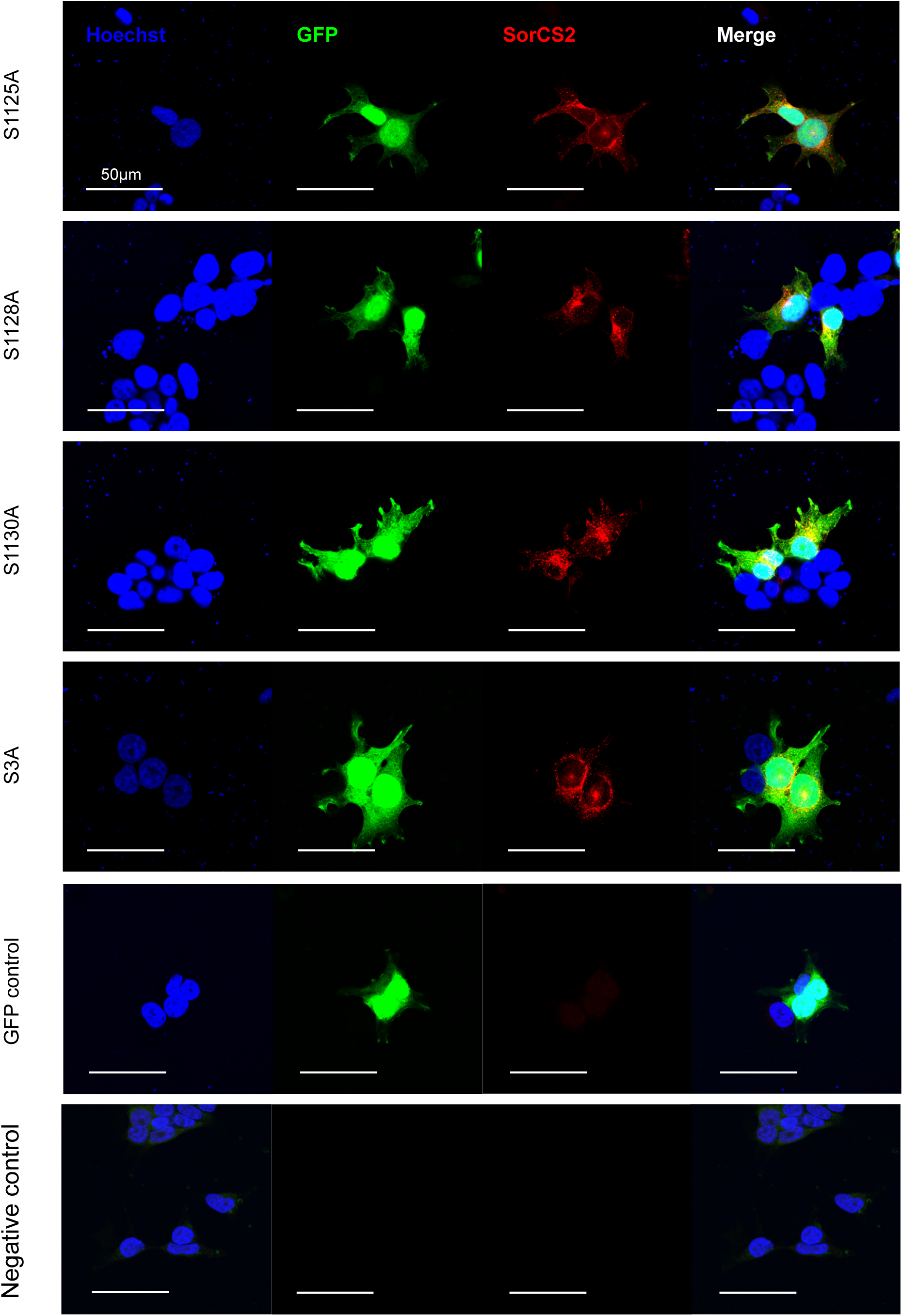
Immunocytochemistry stainings of SorCS2 in transfected HEK293 cells with GFP and SorCS2 variants with serine-to-alanine mutations in tail (S1125A, S1128A, S1130A, S1125A/S1128A/S1130A (denoted S3A)).

**Supplementary Figure 6:**
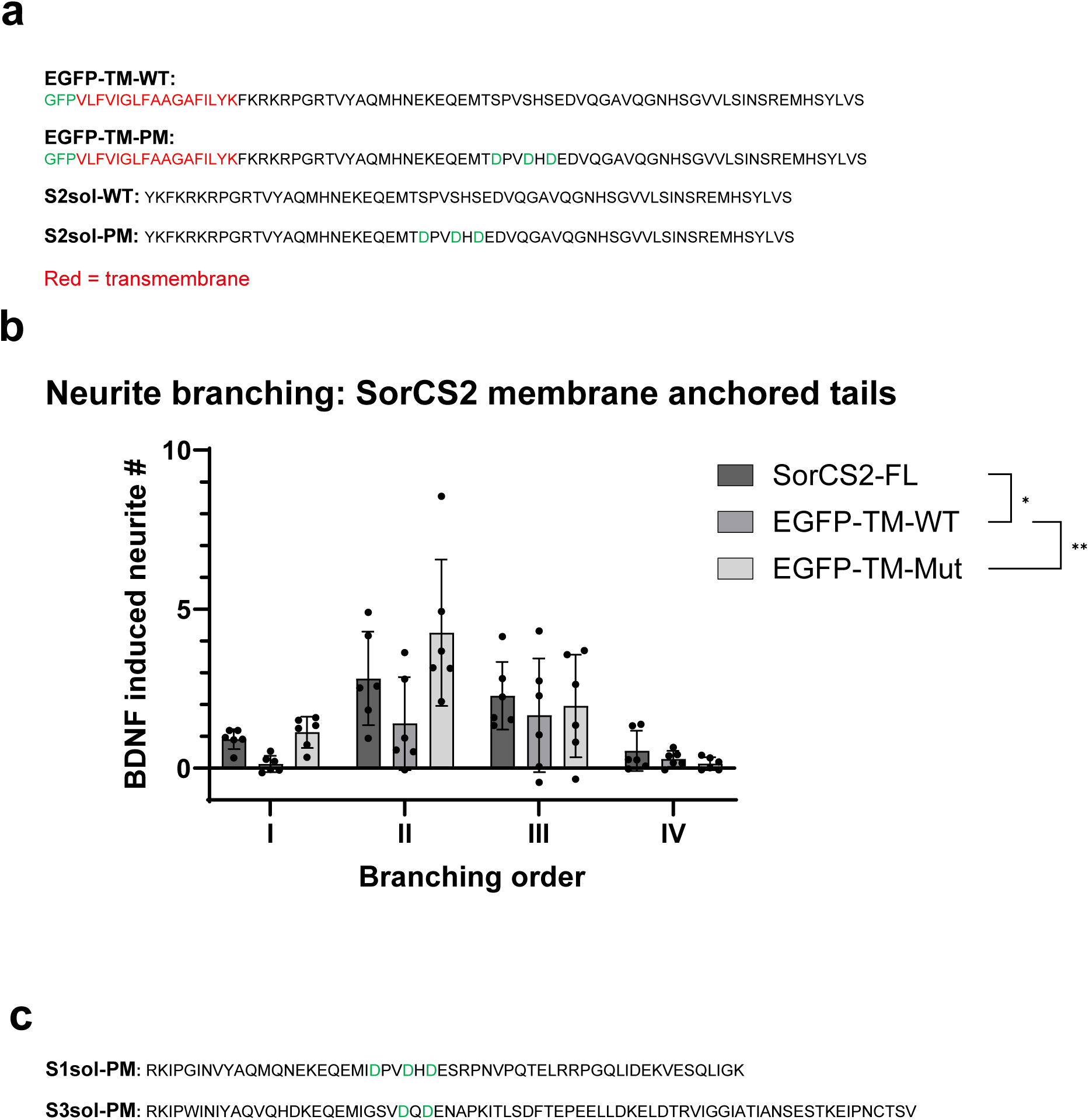
**(a)** Sequences of constructs of both wild-type (WT) and phosphomimetic (PM) SorCS2-tail linked to EGFP and the corresponding free ICDs. **(b)** Extended data on neuronal branching orders in response to BDNF (1nM) in SorCS2 KO hippocampal neurons transfected with GFP and SorCS2-tail constructs linked to EGFP or wild-type SorCS2 (WT) (n=4-6 coverslips per group with 5-15 neurons analysed per coverslip). Significance was calculated using ordinary two-way ANOVA of main effects. **p < 0.01, ***p<0.001 and ****p < 0.0001. Data represented as mean ± SD.. **(c)** Sequences of constructs of phosphomimetic (PM) SorCS1 (S1sol-PM) and SorCS3 (S3sol-PM) soluble-tail constructs.

**Supplementary Figure 7:**
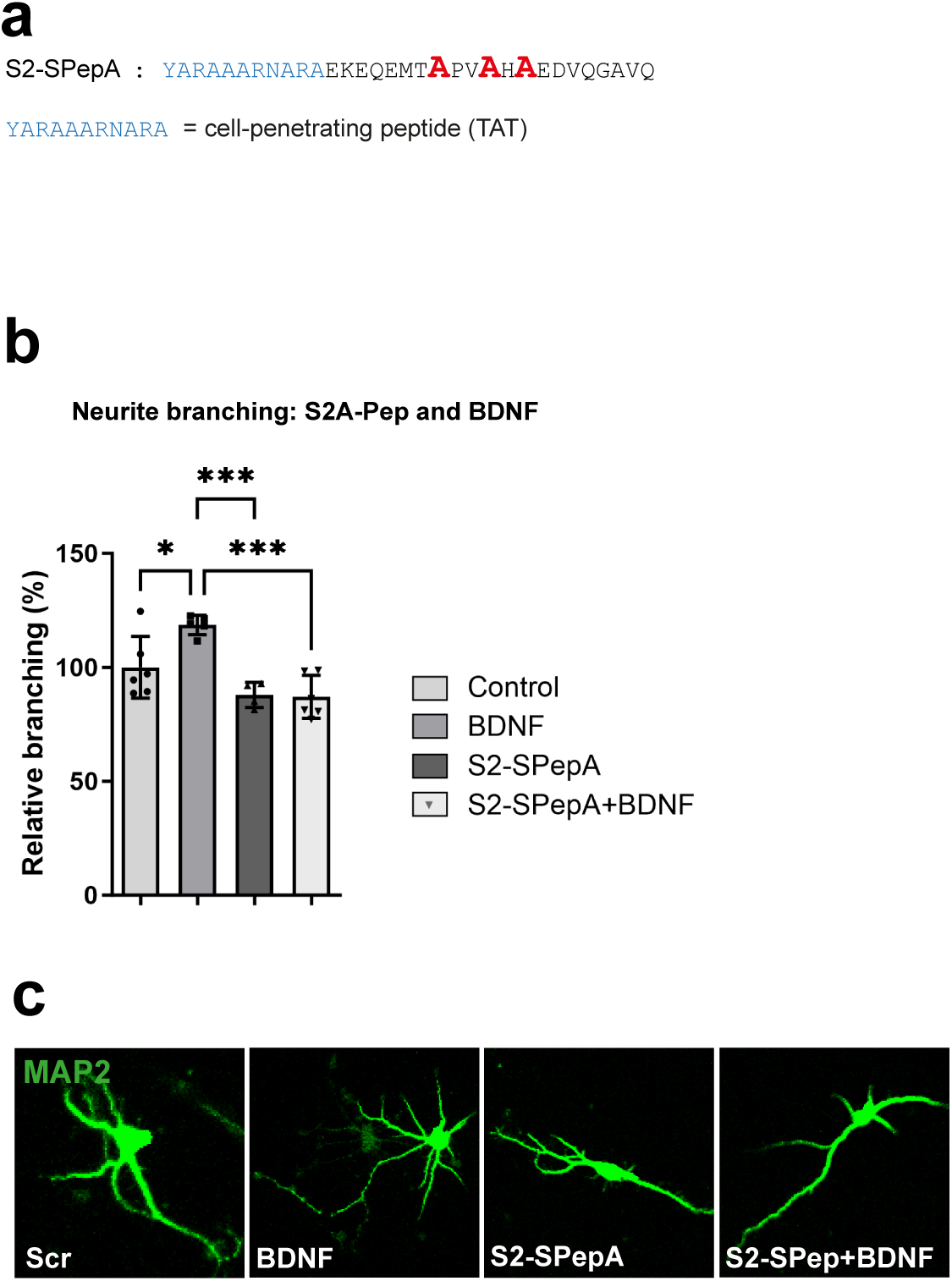
**(a)** Sequences of SorCS2-derived dephospho-peptide. Dephospho-mimetic alanine acids shown in red, TAT-sequence are marked in blue. **(b)** Neurite branching of wild-type (WT) hippocampal neurons treated with outlined compounds and **(c)** representative images (n=4-6 coverslips per group with 15 neurons analysed per coverslip). Significance of neurite branching experiments was calculated using 1-way ANOVA (*p < 0.05). Data represented as mean ± SD.

**Supplementary Figure 8:**
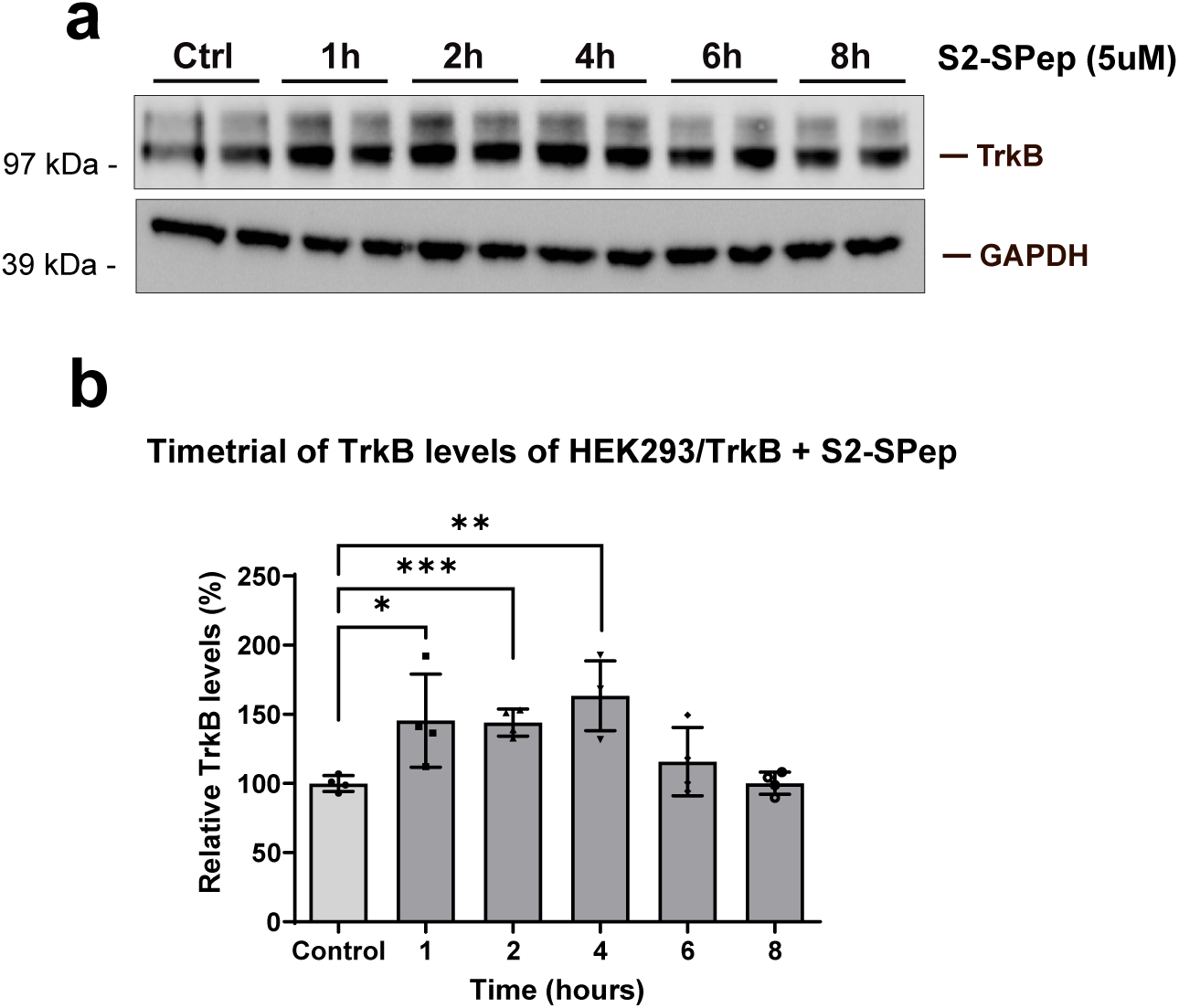
**(a)** Western blot of TrkB levels in HEK293 transfected with TrkB and treated with S2-SPep (5uM) for indicated times and **(b)** densitometric analysis of TrkB relative to GAPDH. Significance was calculated using two-tailed Student’s *t*-test (*p < 0.05, **p < 0.01 and ***p < 0.001). Data represented as mean ± SD.

**Supplementary Figure 9:**
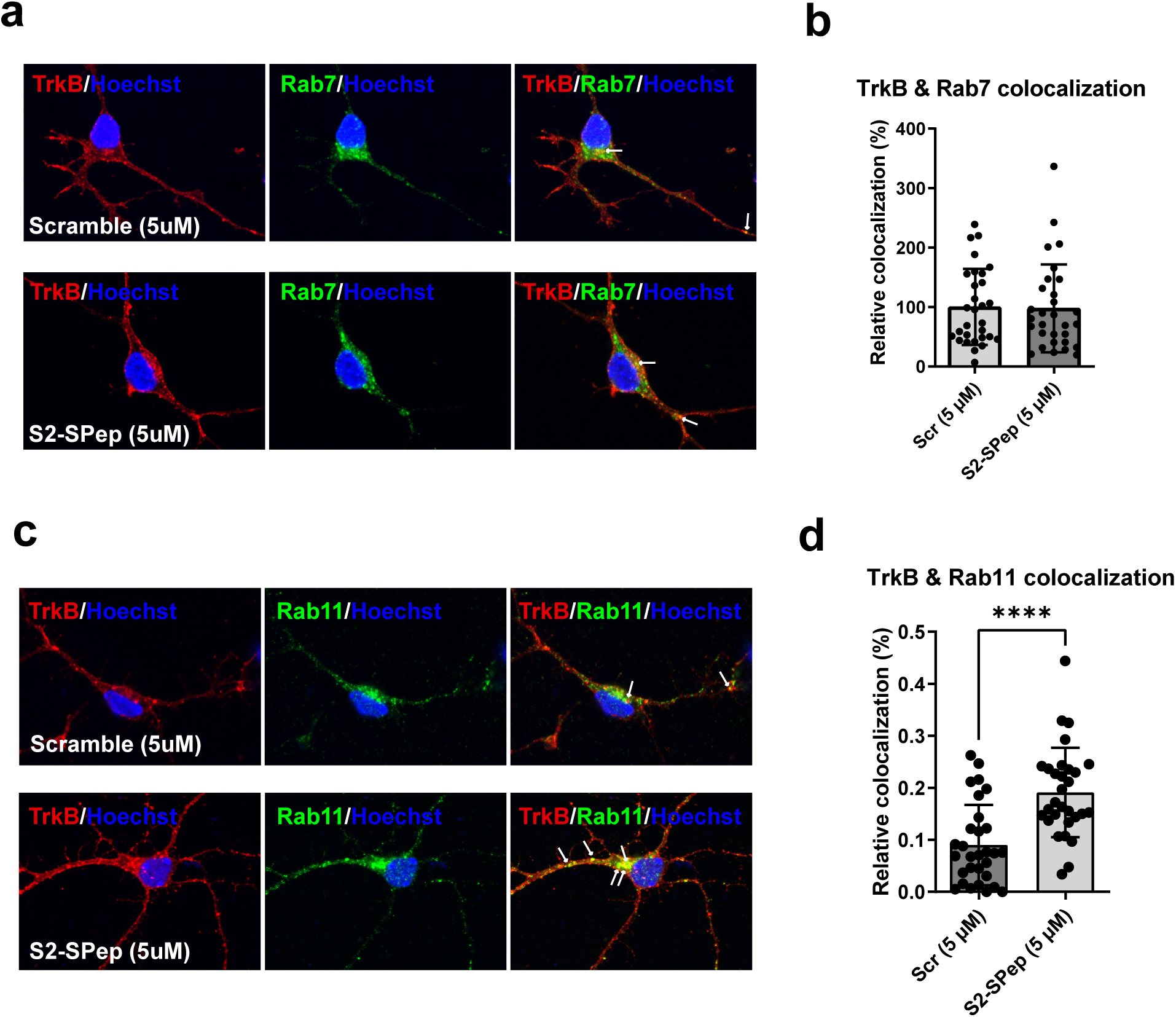
**(a)** Representative confocal images of TrkB and Rab7 following S2-SPep treatment (5uM, 1 hour) and **(b)** the relative weighted colocalization calculated by Zen software. **(c)** Representative confocal images of TrkB and Rab11 following S2-SPep treatment (5uM, 1 hour) and **(d)** the relative weighted colocalization calculated by Zen software. Significance was calculated using two-tailed Student’s *t*-test (*p < 0.05, **p < 0.01 and ***p < 0.001). Data represented as mean ± SD.

**Supplementary Figure 10:**
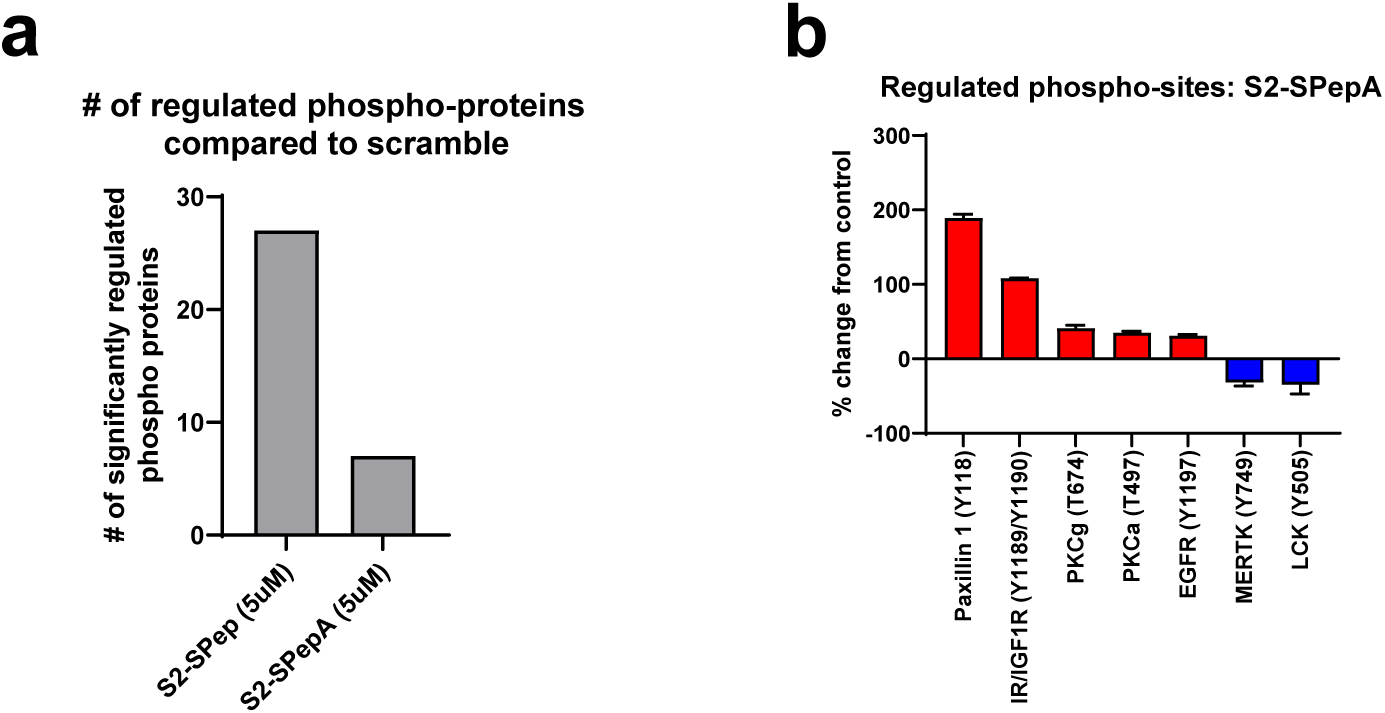
**(a)** Number of identified phospho-proteins significantly regulated by either S2-SPep (5uM, 10min) or S2-SPepA (5uM, 10min) compared to scrambled (5uM, 10min) treated hippocampal neurons using phospho-antibody array. **(b)** Identification of kinase targets of S2-SPepA (5uM, 10 min) relative to Scrambled-treated (5uM, 10 min) wild-type (WT) hippocampal neurons (DIV5) using a phospho-antibody array (n=3). Only significant (p<0.05) targets with 30% change from control and signal intensities above 100 are shown.

**Supplementary Figure 11:**
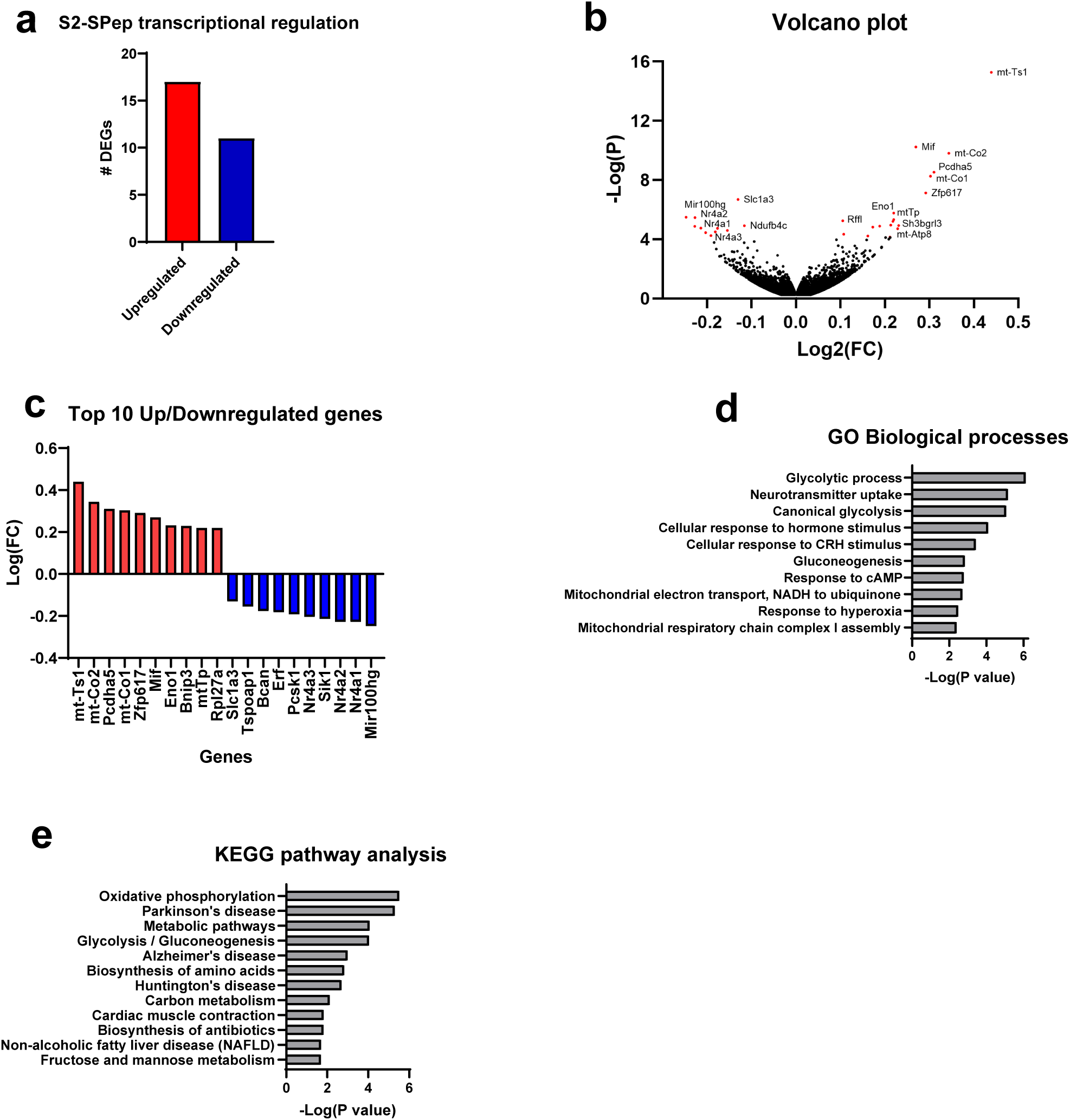
**(a)** Number of differentially expressed genes (DEGs) in wild-type hippocampal neurons induced by S2-SPep (5 µM) relative to Scrambled treated (5 µM) (n=4). **(b)** Volcano plot with DEGs shown in red with annotated gene names. **(c)** Top 10 most upregulated and downregulated genes by S2-SPep treatment. **(d)** Principal component analysis of mRNA sequencing data from wild-type hippocampal neurons treated with S2-SPep (5uM, 2 hours) or Scrambled peptide (5uM, 2 hours) (n=4). **(e)** Enriched GO Biological Processes for all nominally significant genes (p<0.005, n=116 genes). **(f)** Enriched pathways (KEGG pathway analysis) of all nominally significant genes.

**Supplementary Figure 12:**
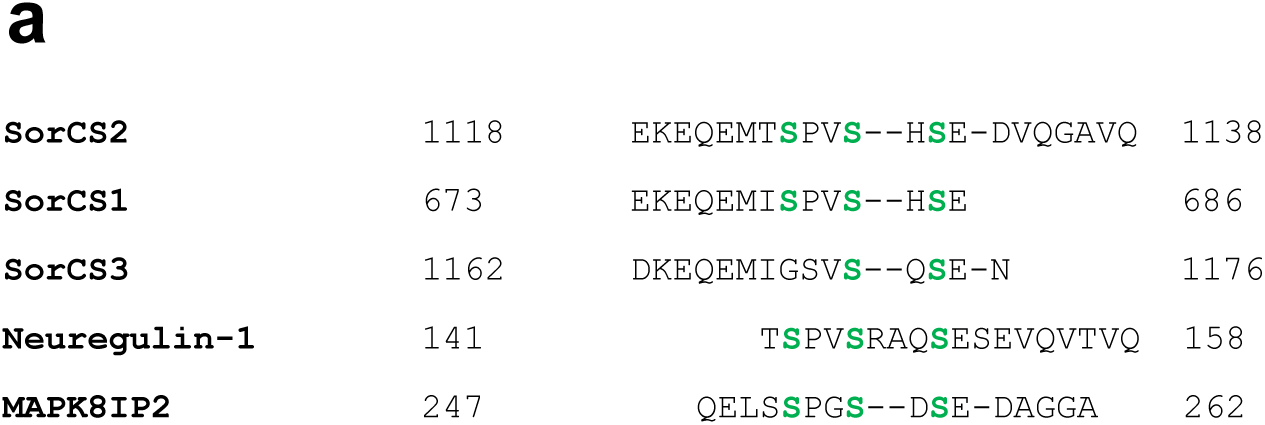
**(a)** Alignment view of top BLAST results from S2-SPep spanning region in SorCS2.

